# The microcephaly protein Abnormal Spindle has an essential role in symmetrically dividing neural precursors to promote brain growth and development

**DOI:** 10.1101/2025.06.27.662046

**Authors:** Shalini Chakraborty, Jack Govaerts, Abigail Hawke, Matthew Werbelow, Todd Schoborg

**Affiliations:** Department of Molecular Biology, University of Wyoming, USA

**Keywords:** Neurogenesis, microcephaly, MCPH, abnormal spindle, ASPM, neuroblasts, neuroepithelium, axon guidance

## Abstract

Neurogenesis must be coordinated in time and space to ensure proper neural development. Defects in this process can lead to a variety of brain malformations that affect tissue size and architecture. Here we examine the spatiotemporal requirements of the Abnormal Spindle (Asp) protein and its human ortholog Abnormal Spindle-Like, Microcephaly Associated (ASPM) in a fly model of human microcephaly (MCPH). By utilizing the fly optic lobe, whose neurogenic program parallels that of mammals, and the genetic tools available in this organism, we uncover the progenitor cell types and critical neurogenesis window necessary for proper brain size and architecture. Asp has an essential function in symmetrically dividing neuroepithelial precursors, yet its expression in asymmetrically dividing neuroblasts alone is not sufficient for optic lobe development. Asp is highly expressed during both embryonic and larval neurogenesis stages, but embryonic expression is dispensable for making a properly sized adult brain. We also show that Asp/ASPM’s interphase nuclear localization is not required for its ability to promote brain development. Expression of the human ASPM N-terminus can also significantly rescue fly brain defects, suggesting conserved mechanisms of function in brain growth control. These results support a model where Asp expression in symmetrically dividing precursors is required to generate a sufficient pool of progenitors prior to the cell fate switch to asymmetrically dividing neuroblasts in order to generate enough neuronal cells to make a properly sized brain.

## Introduction

Neurogenesis is the process by which neural stem cell precursors generate neurons and glia. It therefore plays a key role in defining both the architecture and size of the nervous system. In primates, cell number is the primary ‘scaling factor’ that determines brain size, suggesting that the ability to produce the correct number of neurons in space and time is an essential function of the neurogenesis program to ensure proper brain size (Azevedo et al., 2009; Herculano-Houzel et al., 2007). A reduction in brain size at birth, particularly a head circumference of more than three standard deviations below the population average, is clinically defined as primary microcephaly (MCPH). It is believed that MCPH is the result of defective neurogenesis, leading to the production of fewer neurons and glia during the neurogenic period of development and a smaller brain (Naveed et al., 2018; Thornton and Woods, 2009; Woods and Parker, 2013).

In mammals, neurogenesis is coordinated through the action of distinct neural stem cell populations that can be broadly categorized by their mode of cell division—symmetric and asymmetric. Neuroepithelial cells (NECs) of the early vertebrate neural tube divide symmetrically within the plane of the epithelium to increase the number of progenitors. At the onset of neurogenesis, NECs switch their identity to become asymmetrically dividing radial glial cells (RGCs) of the ventricular zone (VZ), which will generate all the neurons and glia of the cortex. A number of intrinsic and extrinsic cellular factors orchestrate this process, ensuring proper orientation of the mitotic spindle in NECs and RGCs during the appropriate developmental time window (Paridaen and Huttner, 2014). Defects that impair these cell division modes, and ultimately the balance between proliferative symmetric divisions and neurogenic asymmetric divisions, have been suggested to be a major driver of the small brain phenotype seen in MCPH patients (Fish et al., 2006; Jayaraman et al., 2016; Johnson et al., 2018).

*Drosophila melanogaster* and its asymmetrically-dividing neural stem cell populations, known as neuroblasts (NBs), have been a foundational model for elucidating mechanisms of neural stem cell function that are relevant to mammals (Brand and Livesey, 2011; Homem and Knoblich, 2012). More recently, it has become apparent that neurogenesis in the *Drosophila* optic lobe is analogous to neurogenesis in mammals (Contreras et al., 2019; Nériec and Desplan, 2016). For example, the fly optic lobe possesses both symmetrically and asymmetrically dividing neural precursors. The symmetrically dividing neuroepithelial cells (NECs) of the outer proliferation center (OPC) serve to expand the progenitor pool and undergo interkinetic nuclear migration prior to cell division, much like mammalian NECs and RGCs (Bertipaglia et al., 2018; Rujano et al., 2013). Fly NECs also undergo a cell fate switch to asymmetrically dividing medulla neuroblasts (mNBs), which will generate a diverse number of neurons and glia for the visual system. This is analogous to the NEC to RGC fate switch observed in mammals (Paridaen and Huttner, 2014). Furthermore, neuronal and progenitor cell migration has been observed in the fly optic lobe, similar to the migration that mammalian progenitors and neurons undergo to generate the layered architecture of the cortex (Apitz and Salecker, 2015; Chen et al., 2016; Morante et al., 2011; Suzuki et al., 2016).

Mutations in *Abnormal spindle-like, microcephaly associated (ASPM)* are responsible for the majority of human MCPH cases (Létard et al., 2018). The fly ortholog, *abnormal spindle* (*asp*) was originally identified by its mitosis phenotype, consisting of unfocused spindle poles, detached centrosomes, cytokinesis defects, and mitotic delay (Gonzalez et al., 1998; González et al., 1989; Ripoll et al., 1985; Saunders et al., 1997; Wakefield et al., 2001). Asp was later shown to bind to the minus ends of microtubules and localize to spindle poles during metaphase, where it interacts with calmodulin (CaM) to form a ‘molecular glue’ to maintain spindle pole focusing and centrosome-pole cohesion (do Carmo Avides and Glover, 1999; Ito and Goshima, 2015; Schoborg et al., 2015). Asp is also highly expressed in all neural stem cell populations of the larval fly brain, where it localizes to spindle poles during both symmetric and asymmetric division and is associated with the baso-lateral membrane of interphase NECs, although it primarily displays a diffuse cytoplasmic localization during interphase in most neural progenitors (Mannino et al., 2023; Rujano et al., 2013). Previous work showed that Asp and ASPM play a key role in orienting the spindle during mammalian and fly neurogenesis, where it prevented the premature switching of symmetric to asymmetric divisions in NECs. Therefore, it has been suggested that Asp/ASPM plays an essential role in ensuring that the pool of symmetrical dividing NECs is sufficiently expanded prior to the switch to asymmetric divisions by the RGCs in order to ensure enough neurons and glia can be generated during the neurogenic developmental time window, thus ensuring proper brain size (Fish et al., 2006; Rujano et al., 2013). Importantly, disrupted mitotic spindle morphology and centrosome-pole cohesion defects are not the primary drivers of the small brain phenotype per se, at least in *Drosophila* (Mannino et al., 2023; Schoborg et al., 2015).

However, we still lack definitive evidence linking the underlying cell biology defects to the MCPH phenotype in *Asp/ASPM* mutants. This is partly due to the fact that while brain size defects have been reported for *ASPM* mutations in mice and Zebrafish, the size phenotypes are relatively subtle compared to what is observed in humans (Bond et al., 2002; Capecchi and Pozner, 2015; Desir et al., 2008; Fujimori et al., 2014; Jayaraman et al., 2016; Kim et al., 2011; Létard et al., 2018; Passemard et al., 2016; Pulvers et al., 2010; Williams et al., 2015). Ferrets have shown promise as a vertebrate ASPM model for human MCPH, as they share a similar cortex architecture and a 20-40% decrease in brain weight when *ASPM* is mutated (Johnson et al., 2018). However, ferrets lack the genetic tools of traditional model organisms, thus limiting the ability to determine the mechanism of ASPM function(s) in brain growth control.

*Drosophila melanogaster* is a suitable option for overcoming these limitations of vertebrate models of ASPM MCPH. Flies recapitulate the human MCPH phenotype upon loss of *asp* and show a highly significant decrease in overall brain size (∼25%), with a 42% reduction observed specifically in the optic lobes (Rujano et al., 2013; Schoborg et al., 2019, 2015). Furthermore, the extensive suite of genetic tools available in flies, such as the UAS/Gal4 system, allow for the dissection of distinct populations of neuronal cells to the Asp MCPH phenotype (Brand and Perrimon, 1993). This can provide tremendous insight into how Asp coordinates proper brain growth and development spatially, temporally, and potentially in a cell type specific fashion.

We previously showed that the expression of a ∼600 amino acid fragment of Asp’s N-terminus, termed the Asp Minimal Fragment (Asp^MF^) can rescue brain size in *asp* mutants when ubiquitously expressed in the brain. However, cell-type specific analysis using the UAS/Gal4 system showed that expression in neurons and glia was dispensable for brain size, while expression driven by the well-characterized *Inscuteable (Insc)-*Gal4 driver, which is active in most neuroblast and neuron populations of the brain, could restore overall brain size. Intriguingly, region-specific size analysis showed that the central brain was larger in animals expressing Asp^MF^ with *Insc-*Gal4, yet the optic lobes were not completely rescued, with ∼15% reduction in size compared to genotyped-matched control optic lobes. Real time expression and lineage tracing experiments revealed that while *Insc-Gal4* is active in all asymmetrically dividing neuroblasts located medially, including the medulla neuroblasts (mNBs) that will give rise to a significant population of medulla neurons and glia in the adult brain, it is not active in symmetrically dividing neural progenitors, including the neuroepithelial cells (NECs) of the Inner and Outer Proliferation Center (IPC & OPC) and lamina precursor cells (LPCs) (Mannino et al., 2023). This hints at the idea that Asp has an essential role in symmetrically dividing NECs of the optic lobe that ensures proper brain size, although this has not been evaluated to date.

Given these findings and considering that neurogenesis in the fly optic lobe is similar to the process in vertebrates, we decided to evaluate the cell type- and developmental stage-specific expression of Asp in the optic lobe for proper development. Our data indicates that Asp expression during the embryonic neurogenesis period is dispensable for adult optic lobe size and morphology, with expression during the larval neurogenesis period sufficient to restore these phenotypes. We also found that active expression is required in symmetrically dividing neuroepithelial cells (NECs), but not in asymmetrically dividing medulla neuroblasts (mNBs), to rescue optic lobe size and morphology. The N-terminus of the human ASPM ortholog can also significantly rescue these phenotypes when expressed in these cell types, suggesting conservation of function. However, this does not require Asp/ASPM’s nuclear localization. Finally, we also uncover a potential non-cell autonomous role for Asp in promoting optic lobe development. Together, these results highlight an essential role for Asp/ASPM specifically within symmetrically dividing NECs, which may be applicable to humans.

### Materials and Methods Fly stocks and husbandry

All stocks and crosses were maintained on standard cornmeal-agar media at 25°C. The *asp* mutant alleles (*asp^T25^* & *asp^Df^*) and *asp* transgenic rescue lines were previously described (Schoborg et al., 2019, 2015). The ΔNLS transgene strains were made by quick change mutagenesis methods of the Ubi-Asp^FL^ and Ubi-Asp^MF^ plasmids (Agilent, USA). The human ASPM fragment was cloned from a partial cDNA library and inserted into the pPGW Gateway cassette specific for flies. The following GAL4 lines were obtained from the Bloomington Stock Center: *w[1118]; P{w[+mW.hs]=GawB}C855a* (BS#:6990). *P{w[+mW.hs]=GawB}gcm[rA87.C]/CyO* (BS#:35541). *yw, hsFLP;sp-1/cyo; SoxN-GAL4* (kind gift from Xin Li). The G-trace analysis fly strain was *w[*]; P{w[+mC]=UAS-RedStinger}4, P{w[+mC]=UAS-FLP.D}JD1, P{w[+mC]=Ubi-p63E(FRT.STOP)Stinger}9F6/CyO* (BS#: 28280). The endogenously GFP-tagged Asp-fTRG strain (*FlyFos024766(pRedFlp-Hgr)(asp[24519]::2XTY1-SGFP-V5-preTEV-BLRP-3XFLAG)dFRT*) was obtained from the Vienna Stock Center (VS#: 318184). Mutations were verified using Single-wing PCR (Carvalho et al., 2009) and Sanger Sequencing. All UAS- and Ubi-transgenic lines were generated in the *yw* mutant background by BestGene (Chino Hills, CA, USA).

### Developmental Timing

Mated females were placed in egg chambers on grape juice agar plates topped with fresh yeast paste at 25°C. Eggs were collected from the plates within a 20 min laying period and allowed to develop at 25°C for 14-16 hrs to enrich for stage 13-16 embryos. 1^st^ instar larvae were collected from the grape juice agar plates 22-24 hrs after egg laying (AEL). 2^nd^ instar larvae were collected 48-50 hours AEL and were distinguished from the 1^st^ instar by size as well as the presence of the rosette-shaped neuroepithelium in the brain. Early/mid-3^rd^ instar larvae were collected 70-72 hrs AEL. Wandering 3^rd^ instar larvae were collected once they emerged from the food; adult females were collected shortly after eclosion and aged for 3-5 days before dissecting brains.

### Immunohistochemistry & Antibodies

#### Embryos

Embryos were collected from grape juice-agar plates, dechorionated in bleach, fixed and immunostained as described (Diaper and Hirth, 2014). Embryos were then mounted on poly-L-lysine (PLL) coated coverslips and a dollop of silicone grease was added to the four corners to serve as a spacer between the coverslip and the microscope slide. A drop of Aqua-Poly/Mount was then placed in the middle of a microscope slide and the coverslips placed on top and gently pressed to level and spread the mounting media. The slides were then left in a cool, dark chamber for 24-48 hrs before imaging. Antibodies used: anti-FasII (1:50, 1D4, DSHB), anti-Discs large (1:50, 4F3, DSHB), anti-Deadpan (1:100, 11D1BC7, Abcam).

#### First and Second Instar Larva

1^st^ and 2^nd^ instar larvae were collected at desired time points from the egg collection plates and rinsed a few times in water to get rid of any undesired debris. The brains were then roughly dissected, with eye-antennal and wing imaginal discs attached to them in cold 1X PBS. Immunostaining was performed according to the standard protocol for 3^rd^ instar larvae, with a few modifications. 1) 0.1% PBT (1X PBS, Triton-X) was used for all washes. 2) Brains were blocked with 0.1% PBT/5%NGS (1 X PBS, Triton-X, Normal Goat Serum) for 15 mins. 3) Incubation with primary antibody was performed overnight at 4°C. 4) The primary antibody solution was then removed and washed 1×1 min, then 3×20 min in PBT. 5) Secondary antibody incubation was performed for 2-3 hrs on a rotator at room temperature, protected from light. 6) Post-secondary washes were performed as follows: 2 × 15 min. in PBT, then 2 × 15 min in PBS. The slides were then mounted following the Aqua-Poly/Mount procedure for embryos.

#### Third Instar Larva

Brains of the indicated third instar stage were carefully dissected in SF900 media, transferred to 1.5 mL tubes containing 1.2% paraformaldehyde (PFA) diluted in 1xPBS, and fixed for 24 hrs while nutating at 4°C. Fixed brains were then washed in 0.5% PBT for 3×15 min at RT, blocked for 1.5 hrs in 0.5% PBT with 5% normal goat serum, and nutated with the following antibodies: anti-Discs large (1:50, 4F3, DSHB), anti-Elav (1:100, 9F8A9, DSHB), anti-Repo (1:30, 8D12, DSHB), anti-Pros (1:100, MR1A, DSHB), anti-GFP (1:500, AB290, Abcam), anti-Deadpan (1:100, 11D1BC7, Abcam), anti-lamin (1:100, ADL101, DSHB). Brains were nutated for 4 hrs at RT before transferring to 4°C for one overnight. Brains were then washed using 0.5% PBT (4×15 mins.) and incubated 1:500 with secondary antibody (anti-mouse/rabbit Alexa-488/647, Invitrogen/ThermoFisher) for 4 hrs at RT before transferring to 4°C for two overnights. Brains were again washed using 0.5% PBT (4×15 min) and stained with DAPI (0.1 ug/mL) for 5 mins. Following a post-fixation step with 4% PFA at RT for 4 hrs, the brains were washed using 0.5% PBT (4×15 min) and mounted on poly-L-lysine coverslips. The coverslips were transferred through an ethanol dehydration series (30%, 50%, 75%, 95%, 100%, 100%, 100%), cleared using xylene, and mounted in DPX mounting media. Slides were allowed to cure for at least 2 overnights before imaging.

#### Adult Brains

Adults were dipped into 100% EtOH for 3-5 seconds to remove the waxy covering of the cuticle and allow for easier dissection. Brains were dissected in fresh SF900 media and maintained in media for no longer than 15 mins at room temperature prior to fixation. The immunostaining procedure was identical to the 3^rd^ instar larval brain procedure with the following modifications. 1) 2% PFA was used for the initial fixation for 55 mins at RT. 2) The initial blocking was done in 2.5% PBT with 5% normal goat serum. 3) After the initial blocking, the brains were washed 4 times with 0.5% PBT before incubation with primary antibody for two overnights at 4°C. 4) The brains were incubated in secondary antibody for three overnights at 4°C. 5) The primary antibody used for adult brains was anti-Bruchpilot (Brp) (1:30, nc82, DSHB).

#### Eye-antennal discs

Discs from wandering 3^rd^ instar larvae were dissected in fresh SF900 media, then fixed and stained as described elsewhere (Hsiao et al., 2012; Spratford and Kumar, 2014). Antibodies used: anti-Elav (1:100, 9F8A9, DSHB), anti-Repo (1:30, 8D12, DSHB), anti-GFP (1:500, AB290, Abcam). Discs were mounted on poly-L lysine coverslips and placed on a microscope slide containing Aqua Poly/Mount as described above for embryos and larvae.

### Drosophila S2 Cell Culture, Transfection, and Immunostaining

S2 cells were obtained from the *Drosophila* Genome Resource Center S2-DGRC (DGRC Stock 6; https://dgrc.bio.indiana.edu//stock/6; RRID:CVCL_TZ72) and maintained in SF900 insect media containing 1x penicillin/streptomycin at 25°C. Transfection was performed using an Amaxa Nucleofector (Lonza). 2 µg vector was diluted in 100 µl nucleofection solution (50 mM d-mannitol, 15 mM MgCl_2_, 5 mM KCl, and 120 mM NaPO_4_, pH 7.2) and used to resuspend a pellet of ∼4 × 10^6^ cells. This solution was added to a cuvette and electroporated using the S2 cell (G-030) setting. Transfected cells were maintained in six-well plates with 2 ml SF900 at 25°C for 48 hrs before imaging. For immunostaining, transfected cells were allowed to adhere to coverslips coated with 0.5 mg/ml Concavalin A for 60 min in a covered 35-mm dish. Transfected S2 cells were fixed with 4% PFA diluted in PBS for 10 min. Cells were counterstained with DAPI for 1 min and then mounted in Vectashield (Vector Laboratories). Coverslips were sealed using nail polish and imaged immediately.

### Microscopy

All imaging was conducted on an Olympus IX83 microscope fitted with a Yokogawa CSU-W1 Dual Disk SoRa, dual Hamamatsu Orca Flash 4.0 V3 sCMOS cameras, and Plan S-Apo 40x & 100x Si Oil Objectives (NA 1.25, 1.35) operated by cellSens software.

### Microcomputed Tomography (u-CT)

Sample preparation, imaging, and analysis were carried out as previously described (Schoborg, 2020; Schoborg et al., 2019). I2KI was used as the contrast agent. Samples were scanned using a SkyScan 1172 desktop scanner controlled by Skyscan software (Bruker). X-Ray source voltage and current settings: 40 kV, 250 μA, 4 W of power. A cooled 14-bit 11Mp (402×2688) CCD detector coupled to a scintillator was used to collect X-rays converted to photons. Medium camera settings at an image pixel size of 2.95 μm were used for fast scans (∼20 min), which consisted of about 300 projection images. Frame averaging was set to 4. Tomograms were generated using NRecon software (Bruker Micro CT, v1.7.0.4).

### Image Analysis

FIJI (ImageJ, v1.54p) (Schindelin et al., 2012) and IMARIS (Oxford Instruments, v10.2.0) were used to analyze confocal microscopy images. To calculate the volume of the medulla neuroblast region from 3^rd^ instar larval brains, the *Surfaces* tool in IMARIS was used, and the sections were segmented using the *segment manually* option using the α-Dlg and α-Dpn channels to demarcate the central brain from the medulla neuroblast region.

For assessing myosin distribution along the apical-basal NEC axis in FIJI, line scans were generated for five different regions across each neuroepithelium and then averaged. A total of 40 different NE cells (20 each brain lobe) were analyzed for each brain. An average of 10 brains and 400 NE cells were analyzed for each genotype. Due to the differences in apico-basal cell length between cells, only the first 15 pixel intensity values between each region (apical and basal) were used to calculate the average and standard deviation for each neuroepithelium analyzed. The image plane used for visualizing the *Drosophila* OPC neuroepithelium corresponded to a single linear strand of the tissue, rather than the full horseshoe-shaped morphology. The following antibodies were used for the analysis: anti-Myosin Regulatory Light Chain (MRLC), phosphoSer19 (p-MRLC) (1:200, AB3381, Millipore Sigma) and anti-Dlg (1:100, 4F3, DSHB).

For cell length measurements, the line tool from Fiji was used to draw a straight line from the apical to basal end of each NE cell using the anti-Dlg signal as a reference. For each genotype, an average of 100-200 NE cell lengths were measured. The anti-Dlg signal was also used to demarcate the boundaries between adjacent cells. The image plane used for this analysis was the same as used for the p-MRLC measurements.

To calculate adult optic lobe, central brain, and entire brain volumes, Dragonfly (COMET Technologies, v2020.2.0.941) was used. Brains were segmented manually using the ROI painter tool (Schoborg, 2020). For the analysis of the UAS-ASPM^MF^ and *c855a-*Gal4/*Gcm*-Gal4 driver experiments, an in-house AI segmentation model specific for the adult brain, developed with Dragonfly’s AI tool, was used to derive volumes (McDaniel et al., 2025).

### Statistics

Statistical analysis was performed using GraphPad Prism 10 software (v10.5.0, 774).

## Results

### Asp is expressed in all dividing neuronal cell types during the embryonic and larval neurogenesis periods

Optic lobe development begins with the formation of the optic placode during embryogenesis, which then becomes subdivided into two primordia called the inner and outer proliferation centers (IPC and OPC) shortly after larval hatching. The cells of these two regions are collectively referred to as neuroepithelial cells (NECs). They continue to divide symmetrically during the early larval stages to increase the progenitor pool, forming distinct tissue structures (Hakes et al., 2018). NECs are progressively converted into neural progenitors as development proceeds, which generate all neurons and glia of each of the four major visual ganglia. The OPC NECs are converted into medulla neuroblasts (mNBs) by a differentiation wave located medially, while OPC NECs located laterally to the lamina furrow become lamina precursor cells (LPCs). mNBs will divide asymmetrically to give rise to medulla neurons, while LPCs will divide symmetrically to generate lamina neurons after receiving signaling inputs from retina photoreceptors (Apitz and Salecker, 2014). The IPC NECs also exist as subdomains (pIPC, sIPC, dIPC), with distinct populations giving rise to asymmetrically dividing neuroblasts that will generate neurons of the lobula, lobula plate, and inner medulla (Apitz and Salecker, 2015) (Fig. 1A, G).

**Figure 1.**
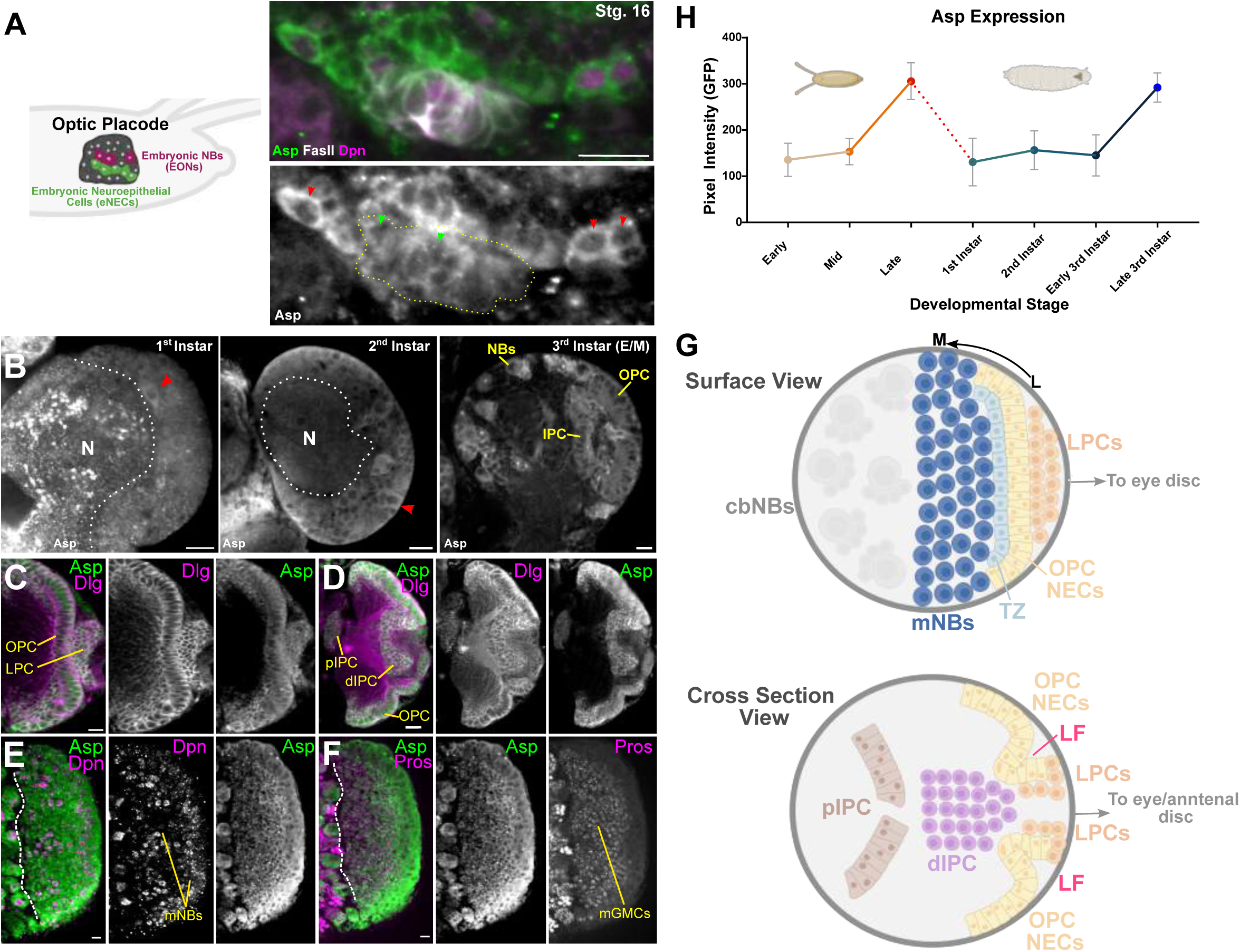
Asp is highly expressed during both the embryonic and larval neurogenesis periods. (A) Asp-FTRG (GFP) expression in the embryonic neuroepithelium (FasII positive) and neuroblasts (EONs, Dpn positive) of a stage 16 embryo. Yellow outline highlights the FasII positive neuroepithelial cells (NECs), green arrow heads mark double FasII and Dpn labeled cells, while red arrowheads mark mature EONs. Asp expression is higher in EONs than in NECs. Note exclusive cytoplasmic localization during interphase. Cartoon shows the relative position of the optic placode in the developing embryo and the relevant cell types. (B) Asp-FTRG (GFP) expression in the first, second, and early/mid (E/M) third instar optic lobe. The neuropil (N) is outlined, Asp-positive neurogenic regions are highlighted by the red arrowheads. Outer Proliferation Center (OPC), Inner Proliferation Center (IPC), Neuroblasts (NBs). (C) Asp-FTRG (GFP) expression in the late third instar optic lobe, showing a more dorsally oriented imaging plane highlighting the OPC and lamina precursor (LPC) regions. Discs large (Dlg) labels cell outlines. (D) A ‘middle’ Z-slice (cross sectional view) of the late third instar larval optic lobe, showing Asp-FTRG (GFP) expression in the OPC and two regions of the IPC, the proximal IPC (pIPC) and the distal IPC (dIPC). (E) Asp-FTRG (GFP) expression in Dpn-positive medulla Neuroblasts (mNBs) of the late third instar optic lobe. White dotted line separates central brain region. (F) Asp-FTRG (GFP) expression in prospero (Pros)-positive ganglion mother cells (mGMCs) derived from the asymmetric division of mNBs. (G) Cartoon representation of the late third instar larval brain when viewed from two different imaging planes (surface and cross section). Relevant cell populations are highlighted. M, medial; L, lateral; mNBs, medulla neuroblasts; transition zone, TZ; Outer proliferation center neuroepithelial cells, OPC NECs; lamina precursor cells, LPCs; central brain neuroblasts, cbNBs; proximal inner proliferation center, pIPC; distal inner proliferation center, dIPC; lamina furrow, LF. (H) Quantification of Asp-FTRG (GFP) levels during early (1-4), mid (5-9), and late (10-17) embryo stages. The dotted line indicates the NB quiescence period. Larval stages as indicated. Scale bars: 10 μm. Cartoons were created using BioRender.

Using a previously validated Asp-GFP line that is expressed under endogenous regulatory elements, we first characterized Asp expression through the embryonic and larval neurogenic periods (Mannino et al., 2023; Sarov et al., 2016). Asp is highly expressed in all dividing neural stem cell populations of the neurogenic embryo, including the embryonic neuroepithelial cells marked by Fasciclin II (FasII) and the embryonic neuroblasts (EONs) marked by Deadpan (Dpn) (Fig. 1A) (Hakes et al., 2018). Furthermore, Asp expression peaks significantly during the late embryonic stages, coinciding with periods of active neurogenesis (Fig. 1H). Following the embryo-larval quiescence period, when neuroblasts resume cell division, Asp expression was observed at lower levels in the first, second, and early/mid third instar larval brain (Fig. 1B, H). However, expression increased dramatically during the late third instar stage when larval neurogenesis is at its highest. Asp is highly expressed in NECs of the OPC and IPC, mNBs, central brain neuroblasts (cbNBs), ganglion mother cells (GMCs) derived from both cbNBs and mNBs, and the lateral LPCs (Fig. 1C-H) (Mannino et al., 2023). Our findings suggest that Asp is expressed in all neuronal precursor cell types that give rise to the entire adult central nervous system (CNS) at both stages of neurogenesis.

### Asp transgenes localize to the interphase nucleus, but this is not required for brain growth and development

In addition to our endogenously expressed Asp-FTRG line, we also generated a series of GFP-tagged Asp transgenes in order to perform genetic rescue assays to better understand Asp’s role in brain growth and development (Schoborg et al., 2019, 2015). We expressed these transgenes using two different promoters: the ubiquitin (ubi-) promoter, which is active in all cell types of the animal, and the UAS promoter to allow for cell- and developmental stage control over Asp transgene expression when paired with the appropriate Gal4 drivers (Brand and Perrimon, 1993). Using the ubi-version of the Asp rescue transgenes, we previously showed that a ∼600 amino acid minimal fragment of Asp’s N-terminus (Asp^MF^) could rescues brain size >90% and tissue morphology defects when expressed ubiquitously in the brain (Mannino et al., 2023; Schoborg et al., 2019), suggesting that this fragment can largely recapitulate the brain growth and development functions of the full length Asp protein.

Analysis of the subcellular localization of the ubi-GFP::Asp^MF^ and ubi-GFP::Asp^FL^ (full length) transgenes revealed a strong interphase nuclear localization for the Asp^MF^ transgene in all cells (Fig 2A-A”,), while Asp^FL^ showed a unique pattern in which a small percentage of cells located adjacent to the lamina furrow showed nuclear localization of GFP::Asp^FL^ during interphase (Fig. 2D-D’). We did not observe this pattern for the full length Asp-FTRG fragment (Fig. S1A), (Mannino et al., 2023), which was exclusively cytoplasmic during interphase in these cell types. Live cell imaging of the ubi-GFP::Asp^FL^ construct revealed that this nucleo-cytoplasmic localization was dynamic, with cells showing nuclear localization of GFP:: Asp^FL^ prior to mitosis, followed by clear cytoplasmic localization in the daughter cells (Fig. 2D”, Video 1). This suggests that Asp’s nuclear localization may be dynamically regulated in a cell type-specific manner.

**Figure 2.**
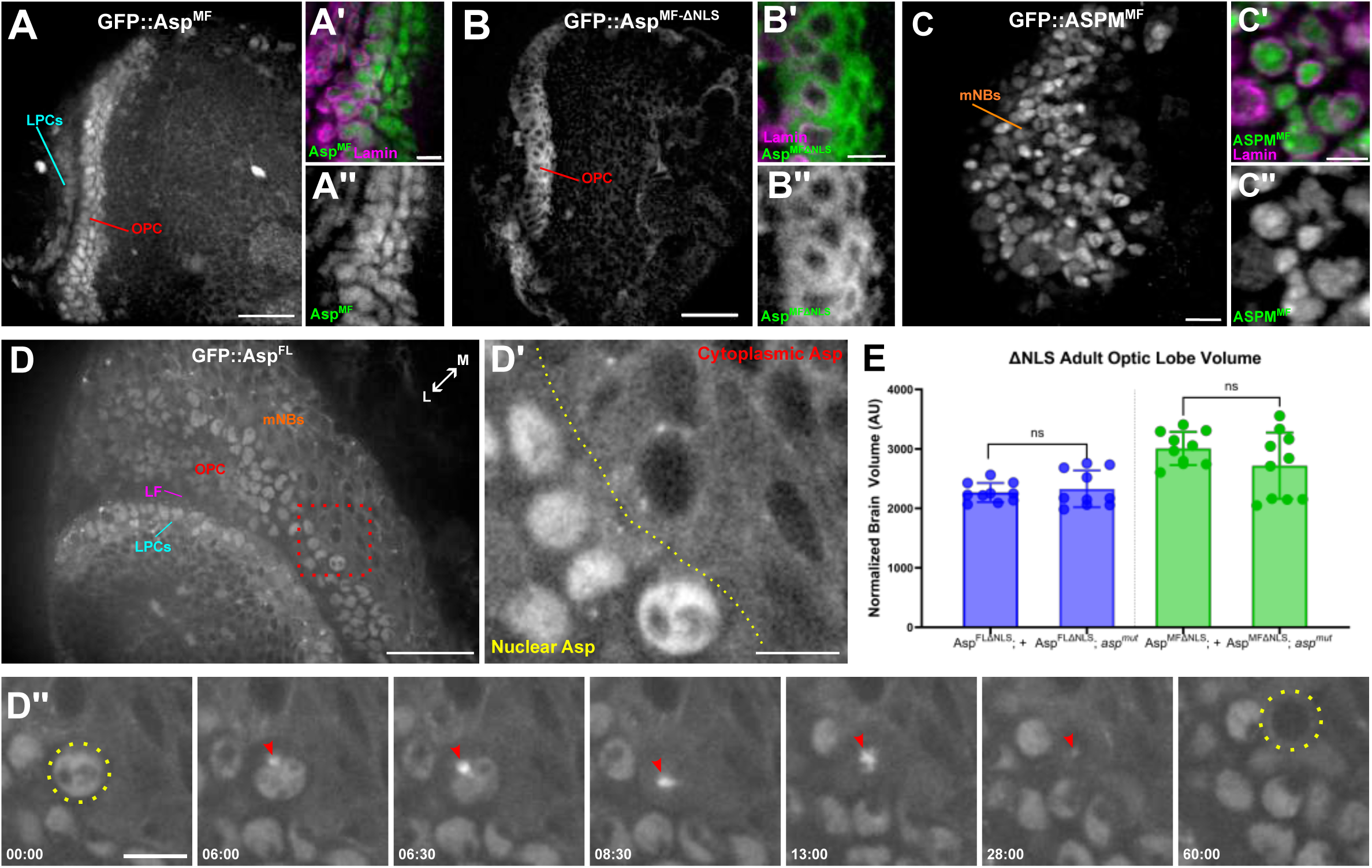
Some Asp transgenes localize to the nucleus of optic lobe stem cell populations during interphase, but this localization is not required for brain growth and development. (A) Localization of the N-terminally tagged Asp^MF^ fragment driven by the ubiquitin (ubi-) promoter in the outer proliferation center neuroepithelial cells (OPC NECs) and lamina precursor cells (LPCs), which is exclusively nuclear as confirmed by nuclear lamin staining (A’, A”). (B) Localization of the N-terminally tagged Asp^MF^ fragment lacking the NLS sequences, driven by the ubi-promoter. OPC NECs are shown. Note localization to the interphase cytoplasm as confirmed by nuclear lamin staining (B, B’). (C) Localization of the N-terminally tagged human ASPM^MF^ fragment showing nuclear localization in the larval brain, confirmed by lamin staining (C’, C”). Medulla neuroblasts (mNBs) are shown, but nuclear localization was also observed in the OPC and LPCs. (D) Live cell imaging still frame of a third instar optic lobe expressing an N-terminally tagged GFP full-length Asp construct driven by the ubi-promoter (ubi-GFP::AspFL). Relevant cell populations and tissue structures are highlighted: medulla neuroblasts (mNBs), neuroepithelial cells of the outer proliferation center (OPC NECs), lamina furrow (LF), lamina precursor cells (LPCs). The red box highlights the region shown in (D’), showing the demarcated zone (yellow line) between nuclear-localized and cytoplasmic Asp. (D”) Still frames from Video 1 showing Asp localized to the interphase nucleus (yellow circle), then transitioning to the astral microtubules radiating out from the apical centrosome during prophase and metaphase (red arrowhead). Following cytokinesis (60:00), Asp no longer shows nuclear localization in one of the daughter cells (yellow circle). Time is shown in minutes:seconds. (E) Genetic rescue assays showing optic lobe volume of ΔNLS Asp^FL^ and Asp^MF^ transgene expression in *asp* mutants (*asp^mut^*) and genotype-matched controls (+). n≥5 brains, Welch’s t-test. ns, P>0.05; *P≤0.05; **P≤0.01; ***P≤0.001; ****P≤0.0001. Error bars represent standard deviation. Medial (M), Lateral (L). Scale bars: (A, B, C) 20 μm, (D) 25 μm, (A’, B’, C’, D”) 5 μm,

Nuclear localization during interphase has been reported for vertebrate ASPM previously (Higgins et al., 2010; Pai et al., 2019; Zhong et al., 2005). We generated humanized flies expressing the orthologous region of fly Asp^MF^, which we termed ASPM^MF^ (a.a. 1-654) (Fig. S1C-D). The human transgene also displayed strong nuclear localization in all cell types of the larval brain in which it was expressed, much like the fly transgene (Fig. 2A-A”, 2C-C”). Using NucPred (Brameier et al., 2007), we computationally identified three NLS sequences in the unstructured region of both fly Asp^MF^ and human ASPM^MF^, although the sequences were not highly conserved (Fig. S1C-D). These three NLS sequences control Asp^MF^’s nuclear localization, as targeted deletion of all three NLS (ΔNLS) sequences was sufficient to exclude nuclear localization of Asp^MF^ in the larval brain and S2 cells (Fig. 2B-B”, Fig. S1C-E).

To determine whether Asp’s nuclear localization is important for fly brain size, we performed a genetic rescue assay using the ΔNLS1-3 version of both Asp^FL^ and Asp^MF^ (Asp^FLΔNLS^ and Asp^MFΔNLS^) expressed ubiquitously in the *asp* mutant (*asp^T25^/asp^Df^)* background and measured adult brain size. Disrupted nuclear localization during interphase did not affect either transgene’s ability to rescue adult optic lobe brain size or other regions of the brain (Fig. 2E, Fig. S5F, F’). Although we cannot rule out that a small pool of nuclear Asp remains upon NLS deletion that is sufficient to promote proper brain growth, given the lack of any observable nuclear localization for the validated Asp-FTRG construct (Fig. S1A), we conclude that Asp’s prominent nuclear localization during interphase is not a primary requirement for proper brain growth and development in flies.

### Assessing the requirements of Asp/ASPM expression in a cell and developmental stage manner for proper development of the fly optic lobe

We next evaluated the spatial and temporal expression requirements for fly Asp and human ASPM in the developing optic lobe that is essential for brain growth and development. We selected three Gal4 drivers that would allow us to dissect the major NEC and Neuroblast populations of the optic lobe at both the embryonic and larval stages: *c855a-*Gal4 (all symmetrically dividing NECs of the outer and inner proliferation centers (OPC and IPC) and the lateral-most lamina precursor cells (LPCs) (Hrdlicka et al., 2002); *Gcm*-Gal4 (lateral-most lamina precursor cells only) (Chotard et al., 2005); and *SoxN-*Gal4, which expresses in asymmetrically dividing medulla neuroblasts (mNBs) (Ray and Li, 2022; Zhu et al., 2022). These drivers were used to express both fly (UAS-GFP::Asp^FL^ and UAS-GFP::Asp^MF^) and human (UAS-GFP::ASPM^MF^) transgenes in *asp* mutant animals (*asp^T25^/asp^Df^)* (Mannino et al., 2023; Schoborg et al., 2019). We then evaluated the ability of each driver and Asp/ASPM transgene to rescue *asp* brain phenotypes, including adult optic lobe size/volume, adult optic lobe neuropil morphology, and larval neuroepithelial morphology (Mannino et al., 2023; Rujano et al., 2013; Schoborg et al., 2019).

We first validated each Gal4 line using the G-TRACE reporter system (Evans et al., 2009). This reporter allows for real-time and lineage tracing analysis of Gal4 activity to determine when and where expression occurs. Since Asp is expressed in all neural precursor cells at both embryonic and larval neurogenesis periods (Fig. 1) (Mannino et al., 2023), we assessed GTRACE reporter activity for each driver throughout the entire neurodevelopmental window (Fig S2A-F,Fig S3A-B”, S4). We also included the late third instar eye/antennal imaginal disc, given the role of photoreceptor cells (R) in the development of the larval optic lobe and lamina and Asp’s ubiquitous expression throughout the tissue, particularly within the morphogenetic furrow (Apitz and Salecker, 2014) (Fig. S3C-E’’’).

In agreement with previous reports, we found that *c855a-*Gal4 is not active in the embryo (Fig. S2A-A’’’, S2D-D’’’) (Hakes et al., 2018), and active expression only becomes evident in late 2^nd^/early 3^rd^ instar in the neuroepithelium (Fig. S3A-A’’). By the late third instar, when neurogenesis is at its peak, *c855a* showed active expression in all symmetrically dividing precursors (OPC, IPC, and LPCs) and lineage expression in asymmetrically dividing medulla neuroblasts (mNBs), medulla ganglion mother cells (GMCs), and medulla neurons (Fig. S4A-A”) (Hrdlicka et al., 2002). In the eye/antennal disc, real time expression was observed in a variegated pattern in the developing eye and antennal fields with ubiquitous lineage expression in all cells, including ommatidial clusters (Fig. S3D-D’’’). Thus, *c855a* is a larval-specific driver active in all symmetrically dividing neural precursors of the larval brain and a subpopulation of eye cells.

*Gcm-*Gal4 showed restricted expression in the embryo, where real time expression in glial cells was evident but not in embryonic neuroblasts or neuroepithelial cells of the optic placode (Fig. S2B-B’’’, S2E-E’’’). This restricted expression to glial cells remained throughout the larval L1 and L2 periods, where we did not find active expression in lamina precursor cells (LPCs) until the late third instar stage (Fig. S3B-B’’, S4B-B’’). While small patches of lineage expression could be seen in parts of the medial OPC, mNBs, and medulla neurons, both real time and lineage expression for *Gcm* were absent from the entirety of the IPC, including all sub-domains (pIPC, dIPC, sIPC, and EMT-like migratory precursors) (Figs. S4B-B’’) (Apitz and Salecker, 2015). This restricted expression pattern was recapitulated in the eye/antennal disc, where we only observed *Gcm-*Gal4 real time expression in glial cells of each ommatidial cluster (Figs. S3E-E’’’). Thus, *Gcm*-Gal4 drives active expression in most glial cells throughout development but is restricted to the population of symmetrically dividing precursors (LPCs) located at the lateral-most margin of the larval brain OPC during the final neurogenesis period, in agreement with previous reports (Chotard et al., 2005). Considering that *Repo-*Gal4 is also active in the same glial populations as *Gcm-*Gal4, and expression of UAS-Asp^FL^ or UAS-Asp^MF^ with *Repo-*Gal4 does not rescue any of the brain phenotypes of *asp* mutants (Mannino et al., 2023), we propose that *Gcm-*Gal4’s activity in subsequent experiments reported here reflects the role of the larval LPC population specifically.

*SoxN-*Gal4 also showed restricted expression in the embryo, with expression in a small number of cells within the optic placode that were FasII-negative, suggesting they were not neuroepithelial cells (Figs. S2C-C’’’, S2F-F’’’). Real time expression during the late third instar stage was restricted to deadpan-positive (dpn+) medulla neuroblasts (mNBs), with weak lineage expression observed in regions of the OPC, LPCs, medulla GMCs, and medulla neurons. Thus, *SoxN* is a driver primarily active in the asymmetrically dividing mNBs population in the larval brain, in agreement with other reports (Fig. S4C-C’’) (Ray and Li, 2022; Zhu et al., 2022).

### Asp expression is required in symmetrically dividing NEC cells to promote proper brain size

We next asked if Asp expression in the subpopulations of neural stem cells determined by *c855a-*Gal4, *Gcm*-Gal4, and *SoxN*-Gal4 could rescue adult brain size in *asp* mutants. Using Microcomputed Tomography (μ-CT) to generate highly precise and accurate 3D volumes (Schoborg, 2020; Schoborg et al., 2019) of the adult optic lobes and central brain (Fig. S1B), we found that *asp* mutants display a 34% decrease in optic lobe volume, in agreement with our previous analysis (Fig. 3A) (Mannino et al., 2023; Schoborg et al., 2019). Expression of UAS-Asp^FL^ and UAS-Asp^MF^ transgenes using either *c855a-*Gal4 or *Gcm-*Gal4 resulted in a significant rescue of adult optic lobe size, with a percent decrease of only 2%-6% observed compared to genotype-matched controls (Fig. 3B, C). Expression of these transgenes using *SoxN*-Gal4 did not rescue adult optic lobe size, with the percent decrease in volume nearly identical to *asp* mutants (27%-32% vs. 34% for *asp* mutants) (Fig. 3A, D). These driver/transgene combinations could not rescue the volume of the entire adult brain or the central brain (Fig. S5A-D, S5A’-D’), in agreement with the restricted spatial expression of these drivers (Fig. S2-S4). Similar results were observed for the third instar larval brain, where we also found that the volume of the medulla neuroblast (mNB) region could be significantly rescued by either *c855a*-Gal4 or *Gcm-*Gal4 (Fig. 3E-G).

**Figure 3.**
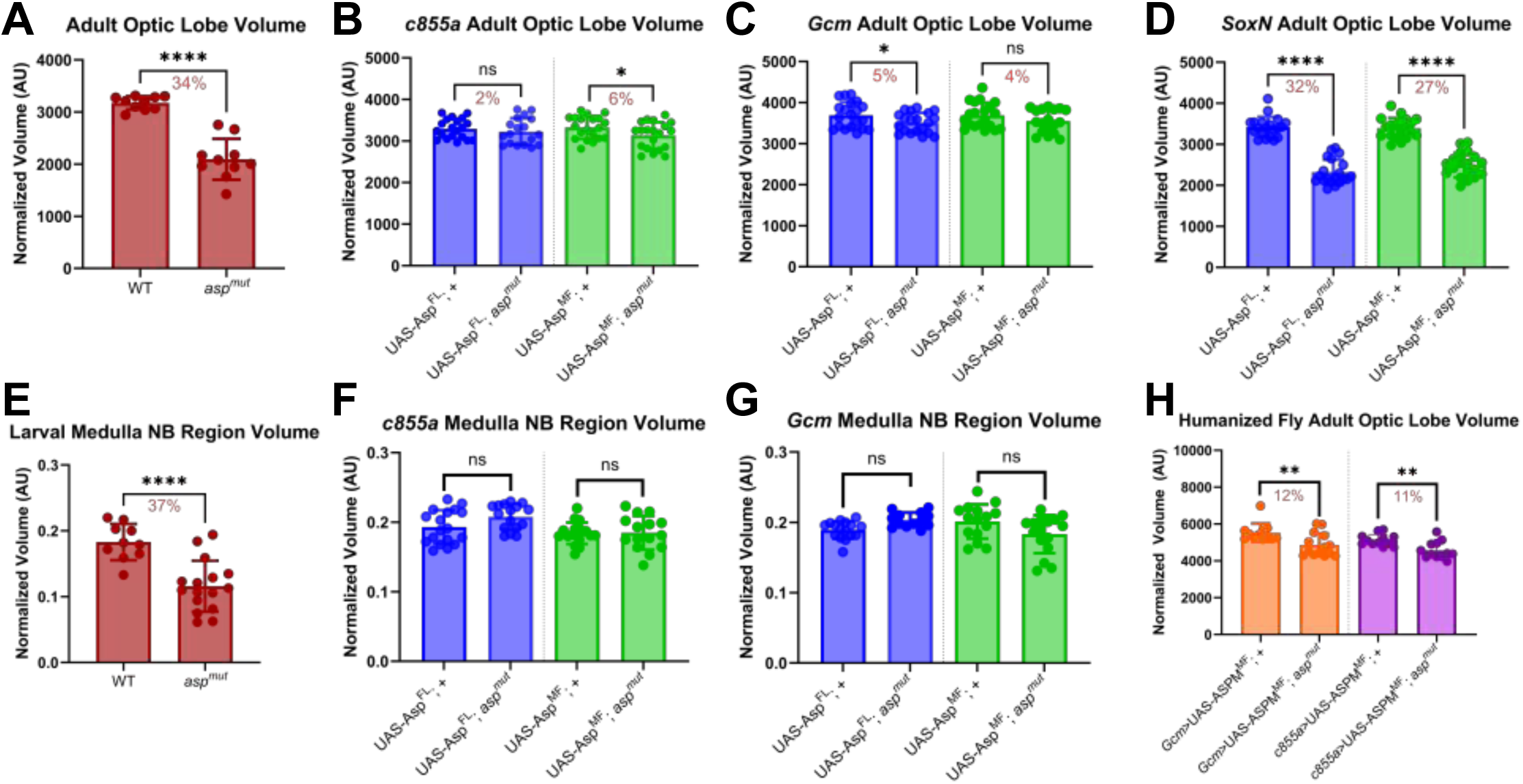
The *asp* microcephaly phenotype can be rescued by expression of Asp transgenes in NECs. Adult optic lobe volumes derived from μ-CT imaging and analysis of *asp* mutants (*asp^mut^*) and genotype-matched controls (+) upon expression of Asp rescue transgenes (UAS-Asp^FL^ and UAS-Asp^MF^) using the indicated Gal4 drivers. (A) Wildtype (WT) and *asp* mutants (*asp^mut^)*, (B) *c855a-*Gal4, (C) *Gcm-*Gal4, (D) *SoxN*-Gal4. Volume of the medulla neuroblast (mNB) region from late third instar larvae of (E) Wildtype (WT) and *asp* mutants (*asp^mut^)*, (F) *c855a-*Gal4, (G) *Gcm-*Gal4. (H) Adult optic lobe volume when human UAS-ASPM^MF^ transgenes are expressed in genotyped-matched controls (+) and *asp* mutants using *c855a-*Gal4 and *Gcm*-Gal4. All volume measurements were normalized to body size (thorax width, adults) or entire optic lobe volume (larvae). Red numbers represent the percent decrease compared to the genotype-matched control. n≥10 brains, Welch’s t-test. ns, P>0.05; *P≤0.05; **P≤0.01; ***P≤0.001; ****P≤0.0001. Error bars represent standard deviation.

We also tested whether our human UAS-ASPM^MF^ transgenes could rescue fly *asp* mutant brain size. Expression with either *c855a*-Gal4 or *Gcm*-Gal4 could significantly rescue optic lobe size, with only an 11-12% reduction in size observed for the optic lobe compared to genotype-matched controls (Fig. 3H). Entire brain and central brain rescue were similar to the percent reduction numbers observed for the fly rescue transgenes (Fig. S5E, E’).

Given the ability of *c855a*- and *Gcm-*Gal4 to significantly rescue optic lobe size when our Asp transgenes were expressed, we wanted to verify that the drivers themselves or the genetic background did not modify the *asp* mutant brain size phenotypes. We confirmed that *c855a*-Gal4 did not modify the *asp* mutant brain size phenotype, particularly in the optic lobe, where a 30% decrease in volume was observed, similar to *asp* mutants alone (Fig. 3A; Fig. S6A-C). On the other hand, *Gcm*-Gal4 on its own could partially suppress the *asp* mutant phenotype, particularly in the optic lobe, where only a ∼10% decrease in size was observed compared to the 34% decrease observed for *asp* mutants (Fig. 3A; Fig. S6D-F). These results suggest that the *Gcm-*Gal4 driver alone, or the genetic background, can influence optic lobe size independently of Asp transgene expression. However, we did observe a better rescue of optic lobe size when *Gcm-*Gal4 was used to express Asp transgenes, with only a 4-5% reduction in size observed for these animals compared to the 10% reduction in size observed with the driver alone (Fig. 3C; Fig. S6D).

These data suggest that the expression of Asp full length or the minimal fragment in symmetrically dividing neuroepithelial cells is sufficient for proper larval and adult optic lobe size. The human ASPM minimal fragment ortholog can also fulfill this role when expressed in fly NECs using these same drivers. Furthermore, given the restricted expression of these NEC drivers to the larval stage, (Fig. S2-S4), our data also suggests that Asp expression in the embryonic stages is dispensable and that larval expression is sufficient to make a properly sized adult optic lobe. Finally, these data suggest that Asp might act non-cell autonomously to promote fly brain growth and development.

### Adult optic lobe neuropil morphology is largely restored by Asp expression in NECs

In addition to the *asp* microcephaly phenotype, *asp* mutant adult brains also show severe morphological defects in the optic lobe, including mispositioning of the neuropils and the cell bodies of the neurons, abnormal projections, disrupted neuropil boundaries, and vacuolar-like structures (Fig. 4A, A’, B, B’). This morphology phenotype appears to be at least partially distinct from the *asp* microcephaly phenotype, considering that certain genetic manipulations can partially restore brain size but not the morphology defects (Mannino et al., 2023; Schoborg et al., 2019). We therefore asked whether the adult optic lobe neuropil morphology could be rescued when our Asp/ASPM transgenes were expressed by *c855a*-Gal4, *Gcm*-Gal4, and *SoxN*-Gal4.

**Figure 4.**
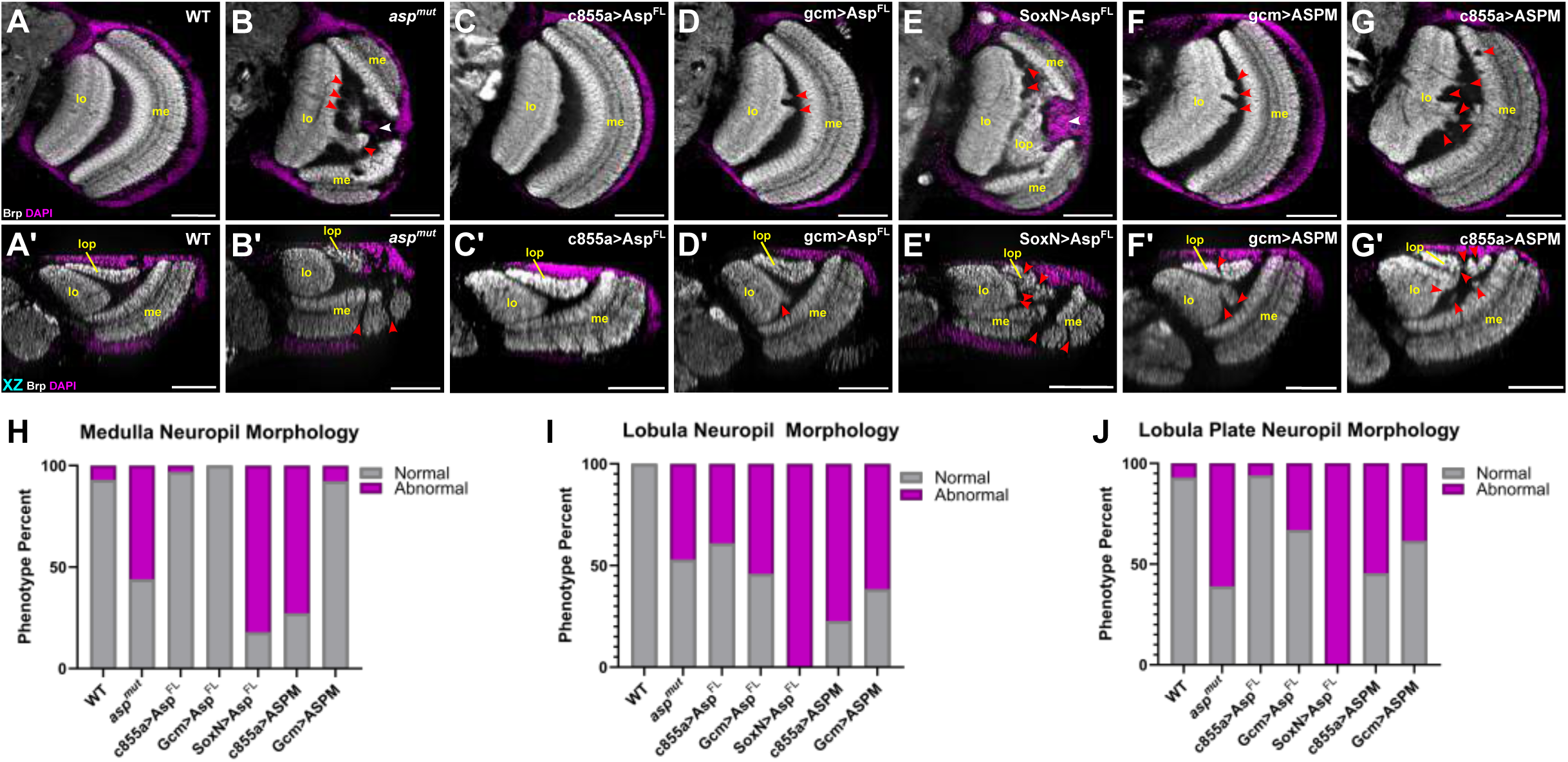
Adult optic lobe neuropil morphology can be rescued by expression of Asp transgenes in NECs. Adult optic lobes stained with α-brp (gray) to visualize the neuropil and DAPI (magenta) to visualize neuronal cell body nuclei upon expression of Asp and ASPM rescue transgenes (UAS-Asp^FL^ and UAS-ASPM^MF^) using the indicated Gal4 drivers. Representative XY (A-G) and XZ (A’-G’) views of the same brain are shown from (A, A’) Wildtype (WT), (B, B’) *asp* mutants (*asp^mut^*), (C, C’) *c855a-*Gal4>UAS-Asp^FL^, (D, D’) *gcm*-Gal4>UAS-Asp^FL^, (E, E’) *SoxN*-Gal4>UAS-Asp^FL^, (F, F’) *gcm*-Gal4>UAS-ASPM^MF^, (G, G’) *c855a*-Gal4>UAS-ASPM^MF^. Quantification of normal vs. abnormal neuropil architecture of the indicated genotypes for the (H) medulla, (I) lobula, (J) lobula plate (n>10 optic lobes). Red arrowheads highlight abnormal neuropil architecture and white arrowheads highlight mispositioned neuronal cell body nuclei. Medulla (me), lobula (lo), lobula plate (lop). Scale bars: 50 μm.

Expression of UAS-Asp^FL^ using the NEC drivers *c855a*-Gal4 and *Gcm*-Gal4 resulted in a significant rescue of the neuropil defects (Fig. 4C, C’, D, D’, H-J). This was most prominent in the medulla neuropil, which showed a complete rescue of morphology in nearly all optic lobes examined (n>10) (Fig. 4H). The lobula and lobula plate rescue was a bit more variable when evaluating the number of brains that had a visible defect in these neuropils (Fig. 4I, J). However, the severity of these defects was much less pronounced compared to the *asp* mutants alone, with all of the observed defects consisting of just minor ‘ruffling’ or small projections emanating from the lobula and lobula plate, with largely intact neuropil boundaries (Fig. 4B, B’, C, C’, D, D’). Similar results were seen for the UAS-Asp^MF^ fragment as well (Fig. S7A, A’, B, B’).

Using *Sox-*Gal4 to express UAS-Asp^FL^ or Asp^MF^ in asymmetrically dividing medulla neuroblasts (mNBs) did not rescue any of the optic lobe neuropil defects compared to *asp* mutants (Fig. 4E, E’, H, I, J; Fig. S7C, C’). Mispositioned neuropils and neuronal cell bodies, abnormal projections, improper boundary formation, and vacuolar structures were evident at a severe level, on par with *asp* mutants alone (Fig. 4B, B’). Finally, we tested whether the expression of the human UAS-ASPM^MF^ transgene with *c855a*-Gal4 and *Gcm*-Gal4 could rescue the neuropil morphology defects. We observed a moderate level of rescue, with phenotypes that were less severe than *asp* mutants alone but not quite to the level seen with the fly transgenes (Fig 4F, F’, G, G’, H, I, J).

Collectively, our data suggests that Asp expression in symmetrically dividing NECs, but not in asymmetrically dividing mNBs, is required for proper adult optic lobe neuropil morphology. This is particularly true for the medulla neuropil. Furthermore, our data suggests that the human ASPM^MF^ fragment can partially recapitulate the function of fly Asp^FL/MF^ in terms of ensuring proper optic lobe neuropil morphology.

### Larval neuroepithelium morphology can be rescued by Asp expression in NECs

Previous studies have shown that Asp contributes to proper neuroepithelium morphology during the larval stage, where it acts via the acto-myosin contractility network to maintain cell shape and tissue organization as a single layer epithelial cell sheet within the OPC (Rujano et al., 2013). We therefore wanted to test whether expression of Asp in our optic lobe cell types of interest could also rescue these phenotypes. We confirmed that our *asp* mutants show a similar neuroepithelial disorganization phenotype in both the OPC and IPC, as reported for other *asp* alleles (Fig. 5A, B, F, G, F’, G’) (Rujano et al., 2013). This phenotype consists of folded, multi-layered, and even rosette-like tissue structures, along with cells that look more rounded with improper spacing/clumping within the epithelial sheet (Fig. 5G, G’). Basally extruded NE cells, a result of dying cells or the loss of cell adhesion, are also visible, particularly within the OPC (Fig. 5G) (Gold and Brand, 2014). The position of the lamina furrow, a visible tissue structure that demarcates the OPC, is also severely disrupted in *asp* mutants (Fig. 5A, B).

**Figure 5.**
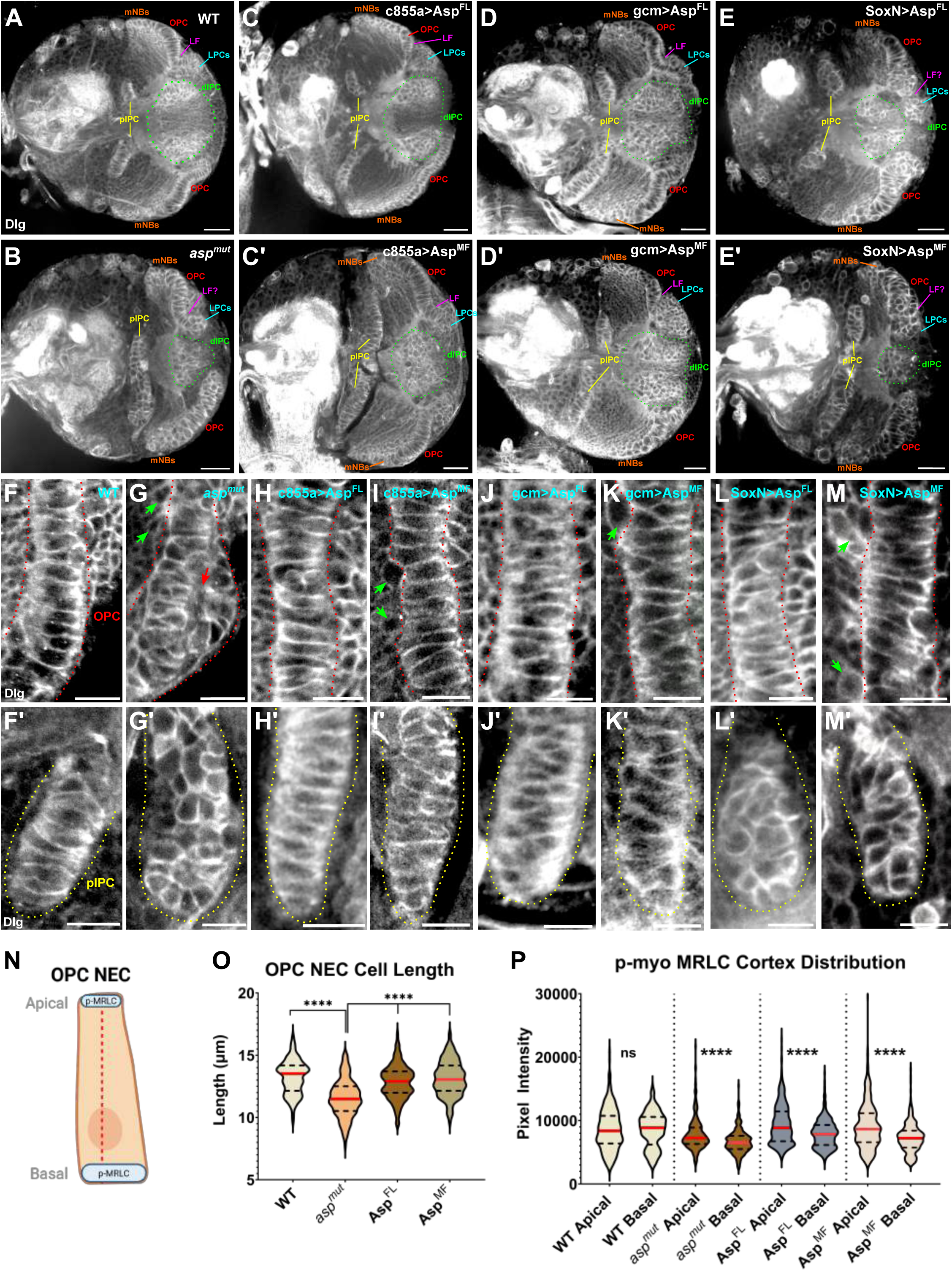
Larval neuroepithelium morphology can be rescued by expression of Asp transgenes in NECs. Representative images of the late third instar larval brain stained with Discs large (Dlg) to visualize cell outlines and neuroepithelium morphology of the Outer Proliferation Center (OPC) and Inner Proliferation Center (IPC) of *asp* mutants upon expression of Asp rescue transgenes (UAS-Asp^FL^ and UAS-Asp^MF^) using the indicated Gal4 drivers. (A) Wildtype (WT), (B) *asp* mutants (*asp^mut^*), (C) *c855a-*Gal4>UAS-Asp^FL^, (C’) *c855a-*Gal4>UAS-Asp^MF^, (D) *gcm*-Gal4>UAS-Asp^FL^, (D’), *gcm*-Gal4>UAS-Asp^MF^, (E) *SoxN*-Gal4>UAS-Asp^FL^, (E’) *SoxN*-Gal4>UAS-Asp^MF^. Close up view of a representative section of the OPC from (F) Wildtype (WT), (G) *asp* mutants (*asp^mut^*), (H) *c855a-*Gal4>UAS-Asp^FL^, (I) *c855a-*Gal4>UAS-Asp^MF^, (J) *gcm*-Gal4>UAS-Asp^FL^, (K), *gcm*-Gal4>UAS-Asp^MF^, (L) *SoxN*-Gal4>UAS-Asp^FL^, (M) *SoxN*-Gal4>UAS-Asp^MF^. Red dashed line demarcates the apical and basal domains of the neuroepithelium. Red arrow indicates rosette-like folds; green arrows highlight basally extruded cells. Shown below each panel is a close-up view of a representative section of the proximal Inner Proliferation Center (IPC) for each genotype (yellow outline) (F’-M’). (N) Cartoon representation of an OPC NEC, highlighting the locations of the active form of the myosin regulatory light chain (p-MRLC) at the apical and basal cortex. Dashed line indicates the position used to derive cell lengths. (O) Cell length measurements (apical-basal) for OPC NECs for the indicated genotypes. (P) Distribution of p-MRLC at the apical and basal domains of OPC NECs for the indicated genotypes. Asp^FL^ and Asp^MF^ transgenes were driven by the *ubi* promoter in (O) and (P). n≥10 brains (∼400 NECs total), Mann-Whitney U Test. ns, ****P≤0.0001. For each violin plot, the red line represents the median, dashed lines represent the interquartile range. Medulla Neuroblasts (mNBs), Outer Proliferation Center (OPC), Lamina furrow (LF), Lamina Precursor Cells (LPCs), distal Inner Proliferation Center (dIPC), proximal Inner Proliferation Center (pIPC). Scale bars: 20 μm (A-E’); 10 μm (F-M’). Cartoon generated using BioRender.

Expression of either Asp^FL^ or Asp^MF^ in both the OPC, IPC, and the lamina precursor cells using *c855a-*Gal4 could significantly rescue this morphology, with the majority of brains showing wildtype morphology (single cell layer packing, columnar cell shape, clearly defined lamina furrow) (Fig. 5C, C’, H, H’, I, I’). There was a higher degree of rescue for Asp^FL^ compared to Asp^MF^ when expressed with this driver, particularly within the IPC (Fig. 5H, H’, I, I’). Neuroepithelial morphology was also restored to a similar degree when Asp^FL^ and Asp^MF^ were expressed only in the lateral-most lamina precursor cells using the *Gcm*-Gal4 driver (Fig. 5D, D’, J, J’, K, K’). However, expression in mNBs using *SoxN*-Gal4 could not restore neuroepithelial morphology in either the OPC or the IPC, with phenotypes nearly identical to that observed in *asp* mutants alone (Fig. 5E, E’, L, L’, M, M’).

In agreement with our qualitative analysis of OPC morphology, we found that the apical to basal distance for individual NEC cells of this region was significantly shorter in *asp* mutants compared to WT animals, suggesting a loss of columnar morphology. Furthermore, expression of the Asp^FL^ or Asp^MF^ rescue transgenes in the *asp* mutant background significantly restored the columnar NEC phenotype (Fig. 5N, O). We also found that the distribution of the active form of the fly myosin regulatory light chain (phosphorylated MRLC) was disrupted in *asp* mutant animals compared to WT, with a higher enrichment at the apical cortex for mutants compared to the basal cortex in WT (Rujano et al., 2013). However, we also found that the p-MRLC distribution did not significantly change upon expression of the Asp^FL^ or Asp^MF^ rescue transgenes, with a lower enrichment at the basal cortex observed, similar to the *asp* mutant alone (Fig. 5P).

Together, these results suggest that larval neuroepithelial tissue architecture can be restored by Asp expression in NECs, but not in neighboring medulla neuroblasts. Also, while we did detect a change in the distribution of the active acto-myosin network between the apical and basal NEC domains, we did not see a significant rescue of this phenotype upon expression of the rescue transgenes in NECs (Rujano et al., 2013), suggesting there may be other cellular processes contributing to the disrupted NEC architecture in *asp* mutants.

### IPC-derived neuroblasts can be restored by expression of Asp in NECs

During our analysis of larval brain tissue morphology, we also noticed that the distal inner proliferation center (dIPC), a horseshoe-shaped region of deadpan-positive (dpn+) and asymmetrically dividing neuroblasts derived from the NECs of the proximal IPC (pIPC) (Apitz and Salecker, 2014), also appeared to be disrupted and showed a significant reduction in overall area/size in *asp* mutants (Fig. 5A, B). We therefore wanted to assess whether this phenotype could be rescued by expression of Asp^FL^ and Asp^MF^ in our cell types of interest. We observed a significant decrease in the number of dpn+ cells within the dIPC of *asp* mutants compared to wildtype (Fig. 6A, A’, B, B’). When either Asp^FL^ or Asp^MF^ were expressed using *c855a*-Gal4 or *Gcm*-Gal4, the population of dpn+ cells was significantly restored to wildtype levels (Fig. 6C, C’, D, D’, E, E’, F, F’). However, expression using *SoxN-*Gal4 could not restore the dpn+ population, with the level of dpn+ signal on par with that observed for *asp* mutants (Fig. 6B, B’, G, G’, H, H’).

**Figure 6.**
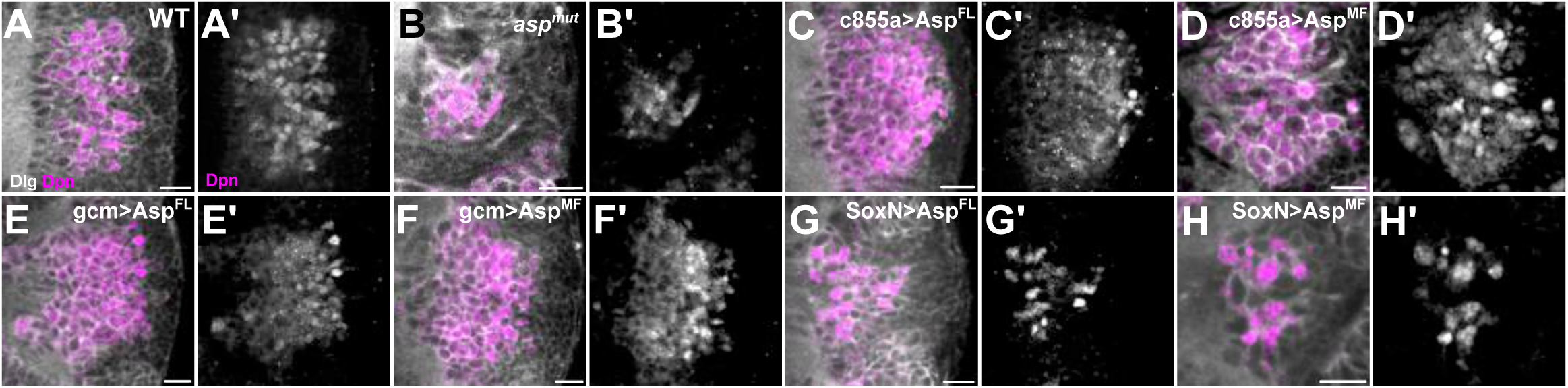
Neuroblast number in the distal IPC is rescued by expression of Asp transgenes in NECs. Representative images of Dpn-positive Neuroblasts (NBs) in the distal inner proliferation center of *asp* mutant larval brains upon expression of Asp rescue transgenes (UAS-Asp^FL^ and UAS-Asp^MF^) using the indicated Gal4 drivers. Discs large (Dlg, gray) and Deadpan (Dpn, magenta) stained images of (A) Wildtype (WT), (B) *asp* mutants (*asp^mut^*), (C) *c855a-*Gal4>UAS-Asp^FL^, (D) *c855a-*Gal4>UAS-Asp^MF^, (E) *gcm*-Gal4>UAS-Asp^FL^, (F), *gcm*-Gal4>UAS-Asp^MF^, (G) *SoxN*-Gal4>UAS-Asp^FL^, (H) *SoxN*-Gal4>UAS-Asp^MF^. The Dpn-only channel is shown in grayscale for the indicating genotypes in A’-H’. Scale bars: 10 μm.

Together, these results suggest that expression of Asp in symmetrically dividing neuroepithelial cells can promote the generation of dpn+ neuroblasts from the proximal IPC (pIPC) in the larval brain, consistent with a model where Asp expression in NECs is required to generate a sufficient pool of precursor cells that will later generate neurons and glia.

## Discussion

The major findings of this investigation are as follows: First, Asp is highly expressed in both symmetric and asymmetric dividing neuronal precursor cells during the embryonic and larval neurogenesis phases in the fly. However, Asp expression is only required during larval neurogenesis to promote proper larval and adult optic lobe development. Embryonic expression is dispensable. Second, active Asp expression during the larval neurogenesis period is required in symmetrically dividing neuroepithelial cells for proper brain growth and tissue architecture of the fly optic lobe. Third, the human ASPM ortholog can largely recapitulate the brain growth and development function of fly Asp, suggesting conservation of function across ∼780 million years of evolutionary divergence. Fourth, Asp’s nuclear localization during interphase is not required for proper brain growth and development. Fifth, active Asp expression in NECs can restore the population of asymmetrically dividing neuroblasts, ensuring a proper neurogenesis program. Finally, Asp may have a non-cell autonomous role in optic lobe development.

Our results show that Asp has an essential role in symmetrically dividing NECs to promote proper brain size, but active expression in asymmetrically dividing neuroblasts is not sufficient. This active Asp expression in NECs could restore a population of asymmetrically dividing neuroblasts. This agrees with a model where the generation of a sufficient pool of neuroepithelial precursors early in neural development is essential prior to the cell fate switch to asymmetrically dividing neural precursors to ensure that the correct number of neurons and glia are generated (Herculano-Houzel et al., 2007; Paridaen and Huttner, 2014) (Fig. 7).

**Figure 7.**
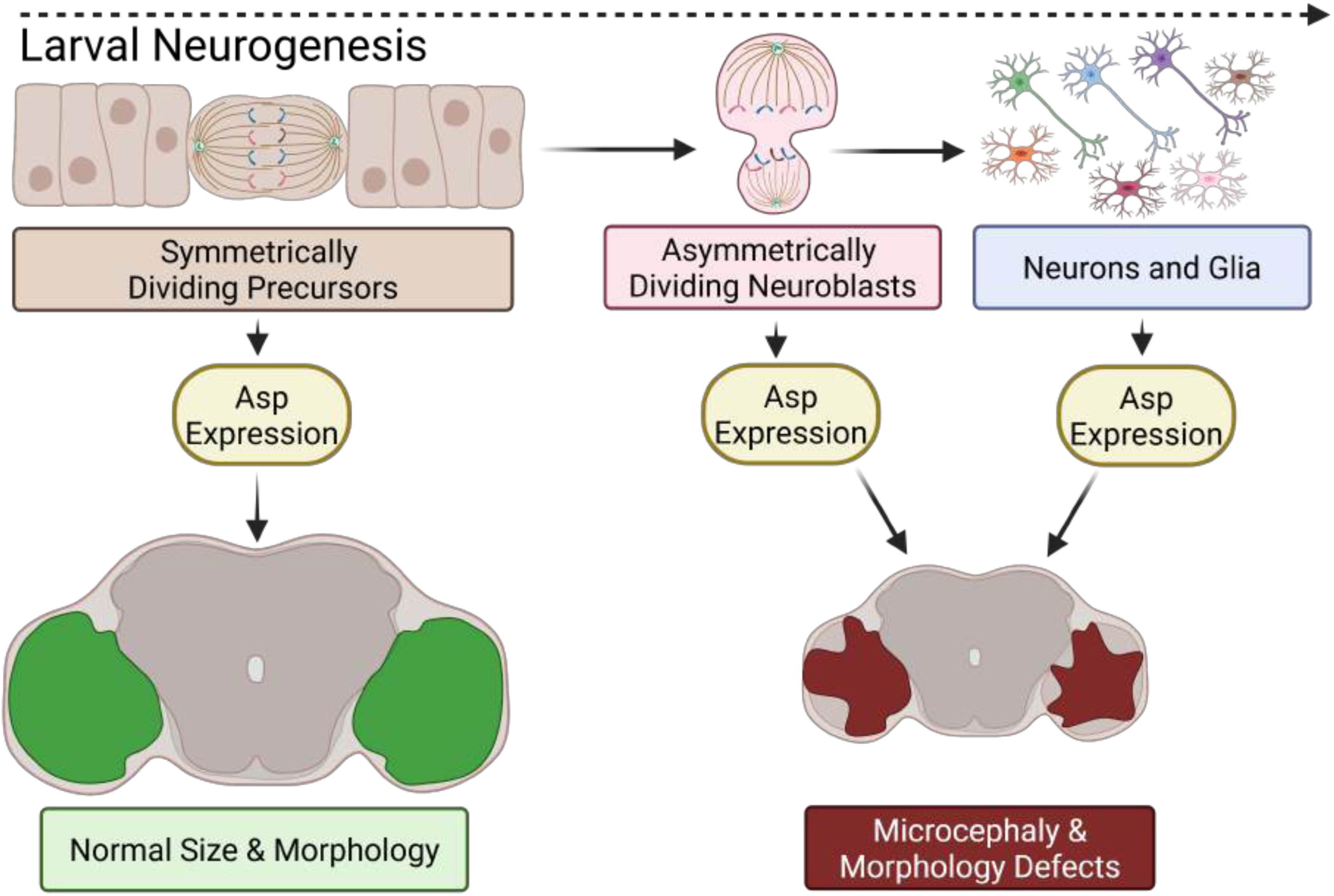
A model for how Asp/ASPM promotes proper optic lobe development. Active expression of Asp during the larval neurogenesis period (third instar) is required in symmetrically dividing neuroepithelial precursors to promote proper optic lobe size and morphology, likely to ensure a sufficient pool of cells prior to the cell fate switch to asymmetrically dividing neuroblasts. Active expression in neuroblasts, neurons, or glia is not sufficient to rescue *asp* microcephaly and morphology defects, as it is likely too ‘late’ in the neurogenesis program to ensure the correct number of neurons and glia to make a properly sized brain. Cartoon created using BioRender.

A key question that remains to be answered is the function of Asp/ASPM in symmetrically dividing NECs to ensure that the quota of progenitors is met. Previous studies in mice showed spindle orientation and cleavage plane defects in symmetrically dividing neuroepithelial cells, which were suggested to alter the pool of symmetrically dividing precursors to asymmetrically dividing neurogenic progenitor cells (Fish et al., 2006). However, studies in flies showed that Asp does not play a significant role in mitotic spindle orientation in neuroblasts (Schoborg et al., 2015), while mutations in other spindle orientation genes, such as mushroom body defect (Mud) and WDR62, do not affect the size of the adult optic lobes (Schoborg et. al, 2019). This suggests there may be other mechanisms that Asp uses to ensure a sufficient pool of neuroepithelial cells. Later mouse studies suggested that ASPM helps to ensure a sufficient pool of neuroepithelial cells by ensuring timely cell cycle progression via regulation of Cdk2/CyclinE activity during G1, ensuring proper self-renewal and maintenance of the NEC progenitor pool (Capecchi and Pozner, 2015). Whether a similar mechanism operates in flies will require further exploration, but our findings highlight two important additional aspects to consider.

First, our humanized flies expressing the orthologous minimal fragment region of human ASPM could significantly rescue adult fly optic lobe size when expressed in NECs, although not to the same degree as the fly minimal fragment. It is possible that there are additional amino acids in the human ASPM protein that were not included in our minimal fragment construct that are required for rescue levels on par with the fly version, and it would be of interest in future studies to assess a full length human ASPM rescue transgene in flies, although this may be difficult given the size of the human coding sequence (>10 Kb for the long isoform). Nonetheless, our results suggest that the molecular function of Asp in NECs is likely conserved in human ASPM, making the fly a relevant model system to understand the etiology of human ASPM microcephaly.

Second, the molecular function of Asp in NECs does not require its nuclear localization during interphase. ASPM has also been shown to localize to the interphase nucleus in various cell types (Higgins et al., 2010; Pai et al., 2019; Zhong et al., 2005). Fly Asp has roles in other tissues, particularly the male and female germline, where we have also observed nuclear localization in a subset of cells. Therefore, we suspect that Asp/ASPM nuclear localization may be essential in these tissues, but not in the brain. Future studies in fly brain NECs should focus on its prominent cytoplasmic localization during interphase for clues regarding its essential molecular function in brain growth control.

Another key finding of our work is that Asp can fulfill its roles as a brain growth regulator during just a single window of neurogenesis in the fly. While it is perhaps not surprising that larval expression of Asp is sufficient to restore adult optic lobe size considering that 90% of adult neurons are generated during this phase (Homem and Knoblich, 2012; Truman and Bate, 1988), our results more broadly suggests the importance of *timing* when it comes to understanding the mechanisms that dictate brain growth and development, regardless of organism. The restricted expression of the *c855a*-Gal4 to primarily the third instar larval period, which lasts for roughly just two days, suggests that the link between developmental time of the organism and the length of neurogenesis are intricately linked to ensure that sufficient neurons and glia are made to ensure proper brain size (Homem and Knoblich, 2012). This is more impressive when considering that we were able to rescue the asymmetrically dividing medulla neuroblast (mNB) region in the late third instar larval brain, considering that this is also the phase in which the OPC NECs are being converted into mNBs (Brand and Livesey, 2011). It suggests that as long as the critical determinants of brain growth such as Asp are functional during brief, but critical windows of neurodevelopment, this is sufficient to ensure proper neurogenesis and subsequent adult optic lobe size. For humans, this would mean simply coordinating similar processes over the span of ∼110 gestation days to generate ∼86 billion neurons, resulting in a properly sized brain (Stepien et al., 2021).

In addition to the *asp* microcephaly phenotype, our findings also explored an even less well-understood aspect of Asp’s neurobiology: morphological defects. Clinically, MCPH is defined as microcephaly vera (true’ microcephaly), where the defining phenotype is simply a small brain but normal tissue architecture. This is to distinguish MCPH from other neurological disorders which may present with additional features such as tissue and skeletal malformations in addition to small brain size (e.g., Sekel syndrome) (Khojah et al., 2021). Mutations in human *ASPM* (MCPH5) are associated with the majority of microcephaly vera cases (Mochida, 2009). However, additional brain tissue malformation may be present in human ASPM patients that have gone undiscovered considering the low spatial resolution of many clinical imaging tools, such as computed tomography (0.5-1 mm) (Lin and Alessio, 2009). More recently, additional brain malformations, such as simplified gyral patterning, have been reported in ASPM patients, suggesting that a lack of imaging resolution in earlier studies may be responsible for the limited understanding of whether ASPM causes morphological defects in human patients (Passemard et al., 2016; Türkyılmaz and Sager, 2021).

In flies, *asp* mutations cause defects in tissue architecture at both the larval and adult stages. Previous studies have shown that Asp can disrupt neuroepithelium architecture in third instar larvae, which was proposed to be due to a disruption in the acto-myosin contractility network within the neuroepithelium due to mislocalization of myosin II between the apical and basal domains (Rujano et al., 2013). Here we showed a nearly identical morphological defect, which could be rescued by expression of Asp transgenes in NECs. However, unlike previous reports, we have not detected an imbalance of myosin II between the apical and basal domains in NECs, calling into question the cellular mechanisms responsible for the larval neuroepithelium phenotype. Loss of neuroepithelium integrity could also be due to a lack of proper adhesion between neighboring cells, leading to the loss of columnar cell morphology and the presence of multiple cell layers (Gold and Brand, 2014). It would be of interest to see if similar adhesion defects are present in the larval brain of *asp* mutants, and if so, the mechanism of how Asp ensures junctional stability.

In adult *asp* mutants, the disrupted neuropil morphology we observe is strikingly similar to the phenotypes observed for boundary formation and axon guidance signaling pathways, such as Netrin and Slit-Robo (Caipo et al., 2020; Suzuki et al., 2018; Tayler et al., 2004). To our knowledge, Asp/ASPM has not been implicated in these pathways, or axon guidance/boundary formation in general. Whether Asp/ASPM is a bona fide regulator of these pathways or plays a more general role in axon guidance via a role at the growth cone, will require further investigation to address this intriguing possibility. In the context of the results shown here, there are three additional considerations. First, expression of the human ASPM minimal fragment could partially rescue morphological defects, suggesting that even if this function is not conserved in humans, the protein itself can partly fulfill this role in flies. Second, why expression in NECs, but not mNBs, is sufficient to rescue the adult morphological defects, particularly of the medulla, will require additional investigation. Third, we are fairly confident that the two *asp* phenotypes (size and morphology) are at least partially independent of one another (Mannino et al., 2023). Yet, both phenotypes can be rescued by the minimal fragment. This suggests that the N-terminus of Asp coordinates multiple biological pathways as a multifunctional (moonlighting) protein to ensure proper brain growth and development.

Finally, our results suggest the possibility that Asp can function in a non-cell autonomous manner to coordinate brain growth and development. This will require additional verification with additional OPC/LPC-only Gal4 drivers or clonal methods, considering the partial suppression we observed for *Gcm*-Gal4 on the *asp* MCPH phenotype alone. However, it is not outside the realm of possibilities, given that ASPM has been recently shown to regulate multiple signaling pathways in several cancer models, such as Wnt, Notch, and Hedgehog (Chan et al., 2024; Cheng et al., 2023; Hsu et al., 2019; Major et al., 2008; Pai et al., 2019; Tsai et al., 2023; Zhang et al., 2021). Furthermore, ASPM was shown to regulate Wnt signaling in the developing mouse brain, where it acts to upregulate Wnt expression, and overactivation of the Wnt pathway alone can rescue several neurogenesis defects in *ASPM* mutants (Buchman et al., 2011).

In flies, Notch plays a key role in virtually all relevant cell types, including NECs, neuroblasts, ganglion mother cells, and neurons (Chen et al., 2016; Contreras et al., 2018; Pinto-Teixeira et al., 2018; Ray and Li, 2022; Sato et al., 2016; Shimojo et al., 2008; Wang et al., 2021, 2011; Yasugi et al., 2010). The fly ortholog of Wnt, Wg, along with the TGF-β member Decapentaplegic (Dpp), has also been shown to promote signaling between distant regions in the fly larval brain to coordinate proper neurogenesis in the IPC (Apitz and Salecker, 2015). Hedgehog also has a well-known role in lamina and optic lobe development via transport along R-cell axons in the eye (Huang and Kunes, 1996). Whether fly Asp serves to regulate these pathways or others (e.g., EGFR) to ensure proper brain growth and development should be prioritized for future studies.

## Acknowledgements

We thank Carey Fagerstrom for cloning of the Asp transgenes, Missy Stuart for fly support, and members of the Schoborg lab for comments and ideas throughout the duration of this project. The *SoxN*-Gal4 driver was a kind gift from Xin Li.

## Data Availability

All raw imaging data can be downloaded from FigShare: https://figshare.com/articles/dataset/All_images_used_for_figures_and_quantification/29428232?file=55734899

## Supplementary Figure Legends

**Supplementary Figure 1. Identification of Asp/ASPM nuclear localization sequences in the minimal fragment region.** (A) Still frame from a live cell imaging experiment of the third instar larval brain expressing the Asp-FTRG full-length construct. Relevant cell populations and tissue structures: medulla neuroblasts (mNBs), outer proliferation center neuroepithelial cells (OPC NECs), lamina precursor cells (LPCs), and lamina furrow (LF). Note lack of interphase nuclear localization for this construct (compare to Fig. 2A). Medial (M), Lateral (L). (B) Cartoon showing the different brain regions (central brain, optic lobe) used for μ-CT volume measurements. Entire brain measurements include both the central brain and the optic lobe. Amino acid sequence of the minimal fragment (MF) regions from both (C) *Drosophila* and (D) human Asp/ASPM. NLS 1, 2, and 3 highlighted in blue. These regions were deleted to make the ΔNLS constructs. (E) NucPred scores and the consequences of deleting each NLS alone or in combination on nuclear localization. Shown are S2 cells transfected with GFP tagged versions of each NLS deletion, co-labelled with DAPI to visualize nuclei. Scale bars: (A) 20 μm, (E) 5 μm.

**Supplementary Figure 2. Embryonic expression patterns of the Gal4 drivers used in this study.** GTRACE analysis in stage 15-16 *Drosophila* embryos. Magenta represents the real time expression pattern; green represents lineage expression. Whole embryo view of (A) *c855a*-Gal4, (B) *gcm*-Gal4, (C) *SoxN*-Gal4 co-labelled with discs large (Dlg) to label cell outlines or deadpan (Dpn) to label embryonic neuroblasts. Insets of the optic lobe placode (outlined by the yellow dashed line) are shown to the right (A’-A’’’; B’-B’’’; C’-C’’’). Insets of the embryonic optic lobe placode stained with the embryonic neuroepithelium marker FasII for (D-D’’’) *c855a*-Gal4, (E-E’’’) *gcm*-Gal4, (F-F’’’) *SoxN*-Gal4. Scale bars: (A-A’, B-B’, C-C’) 20 μm, (D, E, F) 10 μm.

**Supplementary Figure 3. Larval brain and eye-antennal imaginal disc patterns of the Gal4 drivers used in this study.** GTRACE analysis in the first, second, and early third (E3) instar larval brain in (A’-A”) *c855a*-Gal4 and (B-B’’) *gcm*-Gal4. Magenta represents the real time expression pattern; green represents lineage expression. Brains are co-labeled with discs large (Dlg) to visualize cell outlines. (C) Asp-FTRG expression in the eye region of the eye-antennal disc of the late third instar larvae. Arrow highlights the relative position of the morphogenetic furrow. GTRACE analysis in the late third instar eye antennal disc for (D-D’’’) *c855a*-Gal4 and (E-E’’’) *gcm*-Gal4 co-labeled with either Elav (neurons) or Repo (glia). A=anterior, P=posterior. Scale bars: 20 μm.

**Supplementary Figure 4. Late third instar expression patterns of the Gal4 drivers used in this study.** GTRACE analysis in the late third instar larval brain. Magenta represents the real time expression pattern; green represents lineage expression. Brains are co-labeled with discs large (Dlg) to visualize cell outlines. Three different Z-planes (Z#1, Z#2, Z#3) are shown for each driver to highlight expression state and the major cell populations and tissue features: Central Brain Region (CB), medulla Neuroblasts (mNBs), Outer Proliferation Center (OPC). Lamina Furrow (LF), Lamina Precursor Cells (LPCs), proximal Inner Proliferation Center (pIPC), medulla Neurons (mN). (A-A”) *c855a-*Gal4. (B-B”) *Gcm*-Gal4. Asterisks highlight glia cell lineage expression in the mNB region (Z#1). Arrows highlight regions of lineage expression in the mNB and mN regions (Z#1 and Z#2). (C-C”) *SoxN*-Gal4. Scale bars: 20 μm.

**Supplementary Figure 5. Volume measurements of other regions of the adult brain.** Adult entire brain volumes derived from μ-CT imaging and analysis of *asp* mutants (*asp^mut^*) and genotyped-matched controls (+) upon expression of Asp rescue transgenes (UAS-Asp^FL^ and UAS-Asp^MF^) using the indicated Gal4 drivers. (A) Wildtype (WT) and *asp* mutants (*asp^mut^)*, (B) *c855a-*Gal4, (C) *Gcm-*Gal4, (D) *SoxN*-Gal4. (E) Adult entire brain volume when human UAS-ASPM^MF^ transgenes are expressed in genotyped-matched controls (+) and *asp* mutants using *c855a-*Gal4 and *Gcm*-Gal4. (F) Adult entire brain volume with ΔNLS versions of Asp^FL^ and Asp^MF^ expressed ubiquitously in the *asp* mutant background (+ refers to genotyped-matched controls). (A’-F’) Central brain measurements from the indicated genotypes outlined in A-F above. All volume measurements were normalized to body size (thorax width, adults). Red numbers represent the percent decrease compared to the genotyped-matched control. n≥10 brains, Welch’s t-test. ns, P>0.05; *P≤0.05; **P≤0.01; ***P≤0.001; ****P≤0.0001. Error bars represent standard deviation.

**Supplementary Figure 6. Influence of *c855a* and *Gcm* Gal4 drivers on the *asp* mutant microcephaly phenotype.** Volume of the entire brain, central brain, and optic lobe derived from μ-CT imaging of each driver in the *asp* mutant background to determine if they can modify the *asp* mutant brain size phenotype without any UAS rescue transgenes in the genetic background. Volumes of the *c855-*Gal4*; asp^mut^* (A) optic lobes, (B) central brain, (C) entire brain. Volumes of the *Gcm-*Gal4; *asp^mut^* (D) optic lobes, (E) central brain, (F) entire brain. Red numbers indicate the percent decrease in volume compared to the genotyped-matched controls. n≥10 brains, Welch’s t-test. ns, P>0.05; *P≤0.05; **P≤0.01; ***P≤0.001; ****P≤0.0001. Error bars represent standard deviation.

**Supplementary Figure 7. Adult optic lobe neuropil morphology upon expression of Asp^MF^ transgenes.** Adult optic lobes stained with α-brp (gray) to visualize the neuropil and DAPI (magenta) to visualize neuronal cell body nuclei upon expression of UAS-Asp^MF^ rescue transgenes using the indicated Gal4 drivers. Representative XY (A-C) and XZ (A’-C’) views of the same brain are shown from (A, A’) *c855a-*Gal4>UAS-Asp^MF^, (B,B’) *Gcm*-Gal4>UAS-Asp^MF^, (C,C’) *SoxN*-Gal4>UAS-Asp^MF^. Red arrowheads highlight abnormal neuropil architecture. Medulla (me), lobula (lo), lobula plate (lop). Scale bars: 50 μm.

## Video Legends

**Video 1. Ubi-GFP::Asp^FL^ shows differential localization from the nucleus to cytoplasm following cell division in the larval optic lobe.** Full length GFP::Asp construct is nuclear during interphase. Upon entry into mitosis, the signal accumulates at the apical centrosome/spindle pole (∼7 min). Following cytokinesis, the signal is excluded from the nucleus of the daughter cell and remains cytoplasmic. The timestamp is minutes:seconds. Frames were acquired every 30 seconds. Scale bar: 5 μm.

## References

Apitz, H., Salecker, I., 2015. A region-specific neurogenesis mode requires migratory progenitors in the Drosophila visual system. Nat. Neurosci. 18, 46–55. 10.1038/nn.3896.

Apitz, H., Salecker, I., 2014. A challenge of numbers and diversity: neurogenesis in the Drosophila optic lobe. J. Neurogenet. 28, 233–249. 10.3109/01677063.2014.922558.

Azevedo, F.A.C., Carvalho, L.R.B., Grinberg, L.T., Farfel, J.M., Ferretti, R.E.L., Leite, R.E.P., Jacob Filho, W., Lent, R., Herculano-Houzel, S., 2009. Equal numbers of neuronal and nonneuronal cells make the human brain an isometrically scaled-up primate brain. J. Comp. Neurol. 513, 532–541. 10.1002/cne.21974.

Bertipaglia, C., Gonçalves, J.C., Vallee, R.B., 2018. Nuclear migration in mammalian brain development. Semin. Cell Dev. Biol. 82, 57–66. 10.1016/j.semcdb.2017.11.033.

Bond, J., Roberts, E., Mochida, G.H., Hampshire, D.J., Scott, S., Askham, J.M., Springell, K., Mahadevan, M., Crow, Y.J., Markham, A.F., Walsh, C.A., Woods, C.G., 2002. ASPM is a major determinant of cerebral cortical size. Nat. Genet. 32, 316–320. 10.1038/ng995.

Brameier, M., Krings, A., MacCallum, R.M., 2007. NucPred--predicting nuclear localization of proteins. Bioinformatics 23, 1159–1160. 10.1093/bioinformatics/btm066.

Brand, A.H., Livesey, F.J., 2011. Neural stem cell biology in vertebrates and invertebrates: more alike than different? Neuron 70, 719–729. 10.1016/j.neuron.2011.05.016.

Brand, A.H., Perrimon, N., 1993. Targeted gene expression as a means of altering cell fates and generating dominant phenotypes. Development 118, 401–415. 10.1242/dev.118.2.401.

Buchman, J.J., Durak, O., Tsai, L.-H., 2011. ASPM regulates Wnt signaling pathway activity in the developing brain. Genes Dev. 25, 1909–1914. 10.1101/gad.16830211.

Caipo, L., González-Ramírez, M.C., Guzmán-Palma, P., Contreras, E.G., Palominos, T., Fuenzalida-Uribe, N., Hassan, B.A., Campusano, J.M., Sierralta, J., Oliva, C., 2020. Slit neuronal secretion coordinates optic lobe morphogenesis in Drosophila. Dev. Biol. 458, 32–42. 10.1016/j.ydbio.2019.10.004.

Capecchi, M.R., Pozner, A., 2015. ASPM regulates symmetric stem cell division by tuning Cyclin E ubiquitination. Nat. Commun. 6, 8763. 10.1038/ncomms9763.

Carvalho, G.B., Ja, W.W., Benzer, S., 2009. Non-lethal PCR genotyping of single Drosophila. BioTechniques 46, 312–314. 10.2144/000113088.

Chan, T.-S., Cheng, L.-H., Hsu, C.-C., Yang, P.-M., Liao, T.-Y., Hsieh, H.-Y., Lin, P.-C., HuangFu, W.-C., Chuu, C.-P., Tsai, K.K., 2024. ASPM stabilizes the NOTCH intracellular domain 1 and promotes oncogenesis by blocking FBXW7 binding in hepatocellular carcinoma cells. Mol. Oncol. 10.1002/1878-0261.13589.

Cheng, L.-H., Hsu, C.-C., Tsai, H.-W., Liao, W.-Y., Yang, P.-M., Liao, T.-Y., Hsieh, H.-Y., Chan, T.-S., Tsai, K.K., 2023. ASPM activates hedgehog and wnt signaling to promote small cell lung cancer stemness and progression. Cancer Res. 83, 830–844. 10.1158/0008-5472.CAN-22-2496.

Chen, Z., Del Valle Rodriguez, A., Li, X., Erclik, T., Fernandes, V.M., Desplan, C., 2016. A unique class of neural progenitors in the drosophila optic lobe generates both migrating neurons and glia. Cell Rep. 15, 774–786. 10.1016/j.celrep.2016.03.061.

Chotard, C., Leung, W., Salecker, I., 2005. glial cells missing and gcm2 cell autonomously regulate both glial and neuronal development in the visual system of Drosophila. Neuron 48, 237–251. 10.1016/j.neuron.2005.09.019.

Contreras, E.G., Egger, B., Gold, K.S., Brand, A.H., 2018. Dynamic Notch signalling regulates neural stem cell state progression in the Drosophila optic lobe. Neural Dev. 13, 25. 10.1186/s13064-018-0123-8.

Contreras, E.G., Sierralta, J., Oliva, C., 2019. Novel strategies for the generation of neuronal diversity: lessons from the fly visual system. Front. Mol. Neurosci. 12, 140. 10.3389/fnmol.2019.00140.

Desir, J., Cassart, M., David, P., Van Bogaert, P., Abramowicz, M., 2008. Primary microcephaly with ASPM mutation shows simplified cortical gyration with antero-posterior gradient pre- and post-natally. Am. J. Med. Genet. A 146A, 1439–1443. 10.1002/ajmg.a.32312.

Diaper, D.C., Hirth, F., 2014. Immunostaining of the developing embryonic and larval Drosophila brain. Methods Mol. Biol. 1082, 3–17. 10.1007/978-1-62703-655-9_1.

do Carmo Avides, M., Glover, D.M., 1999. Abnormal spindle protein, Asp, and the integrity of mitotic centrosomal microtubule organizing centers. Science 283, 1733–1735. 10.1126/science.283.5408.1733.

Evans, C.J., Olson, J.M., Ngo, K.T., Kim, E., Lee, N.E., Kuoy, E., Patananan, A.N., Sitz, D., Tran, P., Do, M.-T., Yackle, K., Cespedes, A., Hartenstein, V., Call, G.B., Banerjee, U., 2009. G-TRACE: rapid Gal4-based cell lineage analysis in Drosophila. Nat. Methods 6, 603–605. 10.1038/nmeth.1356.

Fish, J.L., Kosodo, Y., Enard, W., Pääbo, S., Huttner, W.B., 2006. Aspm specifically maintains symmetric proliferative divisions of neuroepithelial cells. Proc Natl Acad Sci USA 103, 10438–10443. 10.1073/pnas.0604066103.

Fujimori, A., Itoh, K., Goto, S., Hirakawa, H., Wang, B., Kokubo, T., Kito, S., Tsukamoto, S., Fushiki, S., 2014. Disruption of Aspm causes microcephaly with abnormal neuronal differentiation. Brain Dev. 36, 661–669. 10.1016/j.braindev.2013.10.006.

Gold, K.S., Brand, A.H., 2014. Optix defines a neuroepithelial compartment in the optic lobe of the Drosophila brain. Neural Dev. 9, 18. 10.1186/1749-8104-9-18.

González, C., Molina, I., Casal, J., Ripoll, P., 1989. Gross genetic dissection and interaction of the chromosomal region 95E;96F of Drosophila melanogaster. Genetics 123, 371–377.

Gonzalez, C., Sunkel, C.E., Glover, D.M., 1998. Interactions between mgr, asp, and polo: asp function modulated by polo and needed to maintain the poles of monopolar and bipolar spindles. Chromosoma 107, 452–460. 10.1007/s004120050329.

Hakes, A.E., Otsuki, L., Brand, A.H., 2018. A newly discovered neural stem cell population is generated by the optic lobe neuroepithelium during embryogenesis in Drosophila melanogaster. Development 145. 10.1242/dev.166207.

Herculano-Houzel, S., Collins, C.E., Wong, P., Kaas, J.H., 2007. Cellular scaling rules for primate brains. Proc Natl Acad Sci USA 104, 3562–3567. 10.1073/pnas.0611396104.

Higgins, J., Midgley, C., Bergh, A.-M., Bell, S.M., Askham, J.M., Roberts, E., Binns, R.K., Sharif, S.M., Bennett, C., Glover, D.M., Woods, C.G., Morrison, E.E., Bond, J., 2010. Human ASPM participates in spindle organisation, spindle orientation and cytokinesis. BMC Cell Biol. 11, 85. 10.1186/1471-2121-11-85.

Homem, C.C.F., Knoblich, J.A., 2012. Drosophila neuroblasts: a model for stem cell biology. Development 139, 4297–4310. 10.1242/dev.080515.

Hrdlicka, L., Gibson, M., Kiger, A., Micchelli, C., Schober, M., Schöck, F., Perrimon, N., 2002. Analysis of twenty-four Gal4 lines in Drosophila melanogaster. Genesis 34, 51–57. 10.1002/gene.10125.

Hsiao, H.-Y., Johnston, R.J., Jukam, D., Vasiliauskas, D., Desplan, C., Rister, J., 2012. Dissection and immunohistochemistry of larval, pupal and adult Drosophila retinas. J. Vis. Exp. e4347. 10.3791/4347.

Hsu, C.-C., Liao, W.-Y., Chan, T.-S., Chen, W.-Y., Lee, C.-T., Shan, Y.-S., Huang, P.-J., Hou, Y.-C., Li, C.-R., Tsai, K.K., 2019. The differential distributions of ASPM isoforms and their roles in Wnt signaling, cell cycle progression, and pancreatic cancer prognosis. J. Pathol. 249, 498–508. 10.1002/path.5341.

Huang, Z., Kunes, S., 1996. Hedgehog, transmitted along retinal axons, triggers neurogenesis in the developing visual centers of the Drosophila brain. Cell 86, 411–422. 10.1016/s0092-8674(00)80114-2.

Ito, A., Goshima, G., 2015. Microcephaly protein Asp focuses the minus ends of spindle microtubules at the pole and within the spindle. J. Cell Biol. 211, 999–1009. 10.1083/jcb.201507001.

Jayaraman, D., Kodani, A., Gonzalez, D.M., Mancias, J.D., Mochida, G.H., Vagnoni, C., Johnson, J., Krogan, N., Harper, J.W., Reiter, J.F., Yu, T.W., Bae, B.-I., Walsh, C.A., 2016. Microcephaly proteins wdr62 and aspm define a mother centriole complex regulating centriole biogenesis, apical complex, and cell fate. Neuron 92, 813–828. 10.1016/j.neuron.2016.09.056.

Johnson, M.B., Sun, X., Kodani, A., Borges-Monroy, R., Girskis, K.M., Ryu, S.C., Wang, P.P., Patel, K., Gonzalez, D.M., Woo, Y.M., Yan, Z., Liang, B., Smith, R.S., Chatterjee, M., Coman, D., Papademetris, X., Staib, L.H., Hyder, F., Mandeville, J.B., Grant, P.E., Im, K., Kwak, H., Engelhardt, J.F., Walsh, C.A., Bae, B.-I., 2018. Aspm knockout ferret reveals an evolutionary mechanism governing cerebral cortical size. Nature 556, 370–375. 10.1038/s41586-018-0035-0.

Khojah, O., Alamoudi, S., Aldawsari, N., Babgi, M., Lary, A., 2021. Central nervous system vasculopathy and Seckel syndrome: case illustration and systematic review. Childs Nerv Syst 37, 3847–3860. 10.1007/s00381-021-05284-8.

Kim, H.-T., Lee, M.-S., Choi, J.-H., Jung, J.-Y., Ahn, D.-G., Yeo, S.-Y., Choi, D.-K., Kim, C.-H., 2011. The microcephaly gene aspm is involved in brain development in zebrafish. Biochem. Biophys. Res. Commun. 409, 640–644. 10.1016/j.bbrc.2011.05.056.

Létard, P., Drunat, S., Vial, Y., Duerinckx, S., Ernault, A., Amram, D., Arpin, S., Bertoli, M., Busa, T., Ceulemans, B., Desir, J., Doco-Fenzy, M., Elalaoui, S.C., Devriendt, K., Faivre, L., Francannet, C., Geneviève, D., Gérard, M., Gitiaux, C., Julia, S., Lebon, S., Lubala, T., Mathieu-Dramard, M., Maurey, H., Metreau, J., Nasserereddine, S., Nizon, M., Pierquin, G., Pouvreau, N., Rivier-Ringenbach, C., Rossi, M., Schaefer, E., Sefiani, A., Sigaudy, S., Sznajer, Y., Tunca, Y., Guilmin Crepon, S., Alberti, C., Elmaleh-Bergès, M., Benzacken, B., Wollnick, B., Woods, C.G., Rauch, A., Abramowicz, M., El Ghouzzi, V., Gressens, P., Verloes, A., Passemard, S., 2018. Autosomal recessive primary microcephaly due to ASPM mutations: An update. Hum. Mutat. 39, 319–332. 10.1002/humu.23381.

Lin, E., Alessio, A., 2009. What are the basic concepts of temporal, contrast, and spatial resolution in cardiac CT? J. Cardiovasc. Comput. Tomogr. 3, 403–408. 10.1016/j.jcct.2009.07.003.

Major, M.B., Roberts, B.S., Berndt, J.D., Marine, S., Anastas, J., Chung, N., Ferrer, M., Yi, X., Stoick-Cooper, C.L., von Haller, P.D., Kategaya, L., Chien, A., Angers, S., MacCoss, M., Cleary, M.A., Arthur, W.T., Moon, R.T., 2008. New regulators of Wnt/beta-catenin signaling revealed by integrative molecular screening. Sci. Signal. 1, ra12. 10.1126/scisignal.2000037.

Mannino, Maria C, Cassidy, M.B., Florez, S., Rusan, Z., Chakraborty, S., Schoborg, T., 2023. Mutations in abnormal spindle disrupt temporal transcription factor expression and trigger immune responses in the Drosophila brain. Genetics 225. 10.1093/genetics/iyad188.

McDaniel, J.F., Marsh, M., Schoborg, T., 2025. A deep learning model for accurate segmentation of the Drosophila melanogaster brain from Micro-CT imaging. Dev. Biol. 525, 71–78. 10.1016/j.ydbio.2025.05.027.

Mochida, G.H., 2009. Microcephaly Vera, in: Encyclopedia of Neuroscience. Elsevier, pp. 843–847. 10.1016/B978-008045046-9.01494-7.

Morante, J., Erclik, T., Desplan, C., 2011. Cell migration in Drosophila optic lobe neurons is controlled by eyeless/Pax6. Development 138, 687–693. 10.1242/dev.056069.

Naveed, M., Kazmi, S.K., Amin, M., Asif, Z., Islam, U., Shahid, K., Tehreem, S., 2018. Comprehensive review on the molecular genetics of autosomal recessive primary microcephaly (MCPH). Genet Res (Camb) 100, e7. 10.1017/S0016672318000046.

Nériec, N., Desplan, C., 2016. From the eye to the brain: development of the drosophila visual system. Curr. Top. Dev. Biol. 116, 247–271. 10.1016/bs.ctdb.2015.11.032.

Pai, V.C., Hsu, C.-C., Chan, T.-S., Liao, W.-Y., Chuu, C.-P., Chen, W.-Y., Li, C.-R., Lin, C.-Y., Huang, S.-P., Chen, L.-T., Tsai, K.K., 2019. ASPM promotes prostate cancer stemness and progression by augmenting Wnt-Dvl-3-β-catenin signaling. Oncogene 38, 1340–1353. 10.1038/s41388-018-0497-4.

Paridaen, J.T.M.L., Huttner, W.B., 2014. Neurogenesis during development of the vertebrate central nervous system. EMBO Rep. 15, 351–364. 10.1002/embr.201438447.

Passemard, S., Verloes, A., Billette de Villemeur, T., Boespflug-Tanguy, O., Hernandez, K., Laurent, M., Isidor, B., Alberti, C., Pouvreau, N., Drunat, S., Gérard, B., El Ghouzzi, V., Gallego, J., Elmaleh-Bergès, M., Huttner, W.B., Eliez, S., Gressens, P., Schaer, M., 2016. Abnormal spindle-like microcephaly-associated (ASPM) mutations strongly disrupt neocortical structure but spare the hippocampus and long-term memory. Cortex 74, 158–176. 10.1016/j.cortex.2015.10.010.

Pinto-Teixeira, F., Koo, C., Rossi, A.M., Neriec, N., Bertet, C., Li, X., Del-Valle-Rodriguez, A., Desplan, C., 2018. Development of concurrent retinotopic maps in the fly motion detection circuit. Cell 173, 485–498.e11. 10.1016/j.cell.2018.02.053.

Pulvers, J.N., Bryk, J., Fish, J.L., Wilsch-Bräuninger, M., Arai, Y., Schreier, D., Naumann, R., Helppi, J., Habermann, B., Vogt, J., Nitsch, R., Tóth, A., Enard, W., Pääbo, S., Huttner, W.B., 2010. Mutations in mouse Aspm (abnormal spindle-like microcephaly associated) cause not only microcephaly but also major defects in the germline. Proc Natl Acad Sci USA 107, 16595–16600. 10.1073/pnas.1010494107.

Ray, A., Li, X., 2022. A Notch-dependent transcriptional mechanism controls expression of temporal patterning factors in Drosophila medulla. eLife 11. 10.7554/eLife.75879.

Ripoll, P., Pimpinelli, S., Valdivia, M.M., Avila, J., 1985. A cell division mutant of Drosophila with a functionally abnormal spindle. Cell 41, 907–912. 10.1016/s0092-8674(85)80071-4.

Rujano, M.A., Sanchez-Pulido, L., Pennetier, C., le Dez, G., Basto, R., 2013. The microcephaly protein Asp regulates neuroepithelium morphogenesis by controlling the spatial distribution of myosin II. Nat. Cell Biol. 15, 1294–1306. 10.1038/ncb2858.

Sarov, M., Barz, C., Jambor, H., Hein, M.Y., Schmied, C., Suchold, D., Stender, B., Janosch, S., K J, V.V., Krishnan, R.T., Krishnamoorthy, A., Ferreira, I.R.S., Ejsmont, R.K., Finkl, K., Hasse, S., Kämpfer, P., Plewka, N., Vinis, E., Schloissnig, S., Knust, E., Hartenstein, V., Mann, M., Ramaswami, M., VijayRaghavan, K., Tomancak, P., Schnorrer, F., 2016. A genome-wide resource for the analysis of protein localisation in Drosophila. eLife 5, e12068. 10.7554/eLife.12068.

Sato, M., Yasugi, T., Minami, Y., Miura, T., Nagayama, M., 2016. Notch-mediated lateral inhibition regulates proneural wave propagation when combined with EGF-mediated reaction diffusion. Proc Natl Acad Sci USA 113, E5153–E5162. 10.1073/pnas.1602739113.

Saunders, R.D., Avides, M.C., Howard, T., Gonzalez, C., Glover, D.M., 1997. The Drosophila gene abnormal spindle encodes a novel microtubule-associated protein that associates with the polar regions of the mitotic spindle. J. Cell Biol. 137, 881–890. 10.1083/jcb.137.4.881.

Schindelin, J., Arganda-Carreras, I., Frise, E., Kaynig, V., Longair, M., Pietzsch, T., Preibisch, S., Rueden, C., Saalfeld, S., Schmid, B., Tinevez, J.-Y., White, D.J., Hartenstein, V., Eliceiri, K., Tomancak, P., Cardona, A., 2012. Fiji: an open-source platform for biological-image analysis. Nat. Methods 9, 676–682. 10.1038/nmeth.2019.

Schoborg, T., Zajac, A.L., Fagerstrom, C.J., Guillen, R.X., Rusan, N.M., 2015. An Asp-CaM complex is required for centrosome-pole cohesion and centrosome inheritance in neural stem cells. J. Cell Biol. 211, 987–998. 10.1083/jcb.201509054.

Schoborg, T.A., Smith, S.L., Smith, L.N., Morris, H.D., Rusan, N.M., 2019. Micro-computed tomography as a platform for exploring Drosophila development. Development 146. 10.1242/dev.176685.

Schoborg, T.A., 2020. Whole Animal Imaging of Drosophila melanogaster using Microcomputed Tomography. J. Vis. Exp. 10.3791/61515.

Shimojo, H., Ohtsuka, T., Kageyama, R., 2008. Oscillations in notch signaling regulate maintenance of neural progenitors. Neuron 58, 52–64. 10.1016/j.neuron.2008.02.014.

Spratford, C.M., Kumar, J.P., 2014. Dissection and immunostaining of imaginal discs from Drosophila melanogaster. J. Vis. Exp. 51792. 10.3791/51792.

Stepien, B.K., Vaid, S., Huttner, W.B., 2021. Length of the Neurogenic Period-A Key Determinant for the Generation of Upper-Layer Neurons During Neocortex Development and Evolution. Front. Cell Dev. Biol. 9, 676911. 10.3389/fcell.2021.676911.

Suzuki, T., Hasegawa, E., Nakai, Y., Kaido, M., Takayama, R., Sato, M., 2016. Formation of Neuronal Circuits by Interactions between Neuronal Populations Derived from Different Origins in the Drosophila Visual Center. Cell Rep. 15, 499–509. 10.1016/j.celrep.2016.03.056.

Suzuki, Takumi, Liu, C., Kato, S., Nishimura, K., Takechi, H., Yasugi, T., Takayama, R., Hakeda-Suzuki, S., Suzuki, Takashi, Sato, M., 2018. Netrin signaling defines the regional border in the drosophila visual center. iScience 8, 148–160. 10.1016/j.isci.2018.09.021.

Tayler, T.D., Robichaux, M.B., Garrity, P.A., 2004. Compartmentalization of visual centers in the Drosophila brain requires Slit and Robo proteins. Development 131, 5935–5945. 10.1242/dev.01465.

Thornton, G.K., Woods, C.G., 2009. Primary microcephaly: do all roads lead to Rome? Trends Genet. 25, 501–510. 10.1016/j.tig.2009.09.011.

Truman, J.W., Bate, M., 1988. Spatial and temporal patterns of neurogenesis in the central nervous system of Drosophila melanogaster. Dev. Biol. 125, 145–157. 10.1016/0012-1606(88)90067-X.

Tsai, K.K., Bae, B.-I., Hsu, C.-C., Cheng, L.-H., Shaked, Y., 2023. Oncogenic ASPM is a regulatory hub of developmental and stemness signaling in cancers. Cancer Res. 83, 2993– 3000. 10.1158/0008-5472.CAN-23-0158.

Türkyılmaz, A., Sager, S.G., 2021. Two New Cases of Primary Microcephaly with Neuronal Migration Defect Caused by Truncating Mutations in the *ASPM* Gene. Mol. Syndromol. 1–8. 10.1159/000516201.

Wakefield, J.G., Bonaccorsi, S., Gatti, M., 2001. The drosophila protein asp is involved in microtubule organization during spindle formation and cytokinesis. J. Cell Biol. 153, 637–648. 10.1083/jcb.153.4.637.

Wang, M., Han, X., Liu, C., Takayama, R., Yasugi, T., Ei, S.-I., Nagayama, M., Tanaka, Y., Sato, M., 2021. Intracellular trafficking of Notch orchestrates temporal dynamics of Notch activity in the fly brain. Nat. Commun. 12, 2083. 10.1038/s41467-021-22442-3.

Wang, W., Liu, W., Wang, Y., Zhou, L., Tang, X., Luo, H., 2011. Notch signaling regulates neuroepithelial stem cell maintenance and neuroblast formation in Drosophila optic lobe development. Dev. Biol. 350, 414–428. 10.1016/j.ydbio.2010.12.002.

Williams, S.E., Garcia, I., Crowther, A.J., Li, S., Stewart, A., Liu, H., Lough, K.J., O’Neill, S., Veleta, K., Oyarzabal, E.A., Merrill, J.R., Shih, Y.-Y.I., Gershon, T.R., 2015. Aspm sustains postnatal cerebellar neurogenesis and medulloblastoma growth in mice. Development 142, 3921–3932. 10.1242/dev.124271.

Woods, C.G., Parker, A., 2013. Investigating microcephaly. Arch. Dis. Child. 98, 707–713. 10.1136/archdischild-2012-302882.

Yasugi, T., Sugie, A., Umetsu, D., Tabata, T., 2010. Coordinated sequential action of EGFR and Notch signaling pathways regulates proneural wave progression in the Drosophila optic lobe. Development 137, 3193–3203. 10.1242/dev.048058.

Zhang, H., Yang, X., Zhu, L., Li, Z., Zuo, P., Wang, P., Feng, J., Mi, Y., Zhang, C., Xu, Y., Jin, G., Zhang, J., Ye, H., 2021. ASPM promotes hepatocellular carcinoma progression by activating Wnt/β-catenin signaling through antagonizing autophagy-mediated Dvl2 degradation. FEBS Open Bio 11, 2784–2799. 10.1002/2211-5463.13278.

Zhong, X., Liu, L., Zhao, A., Pfeifer, G.P., Xu, X., 2005. The abnormal spindle-like, microcephaly-associated (ASPM) gene encodes a centrosomal protein. Cell Cycle 4, 1227– 1229. 10.4161/cc.4.9.2029.

Zhu, H., Zhao, S.D., Ray, A., Zhang, Y., Li, X., 2022. A comprehensive temporal patterning gene network in Drosophila medulla neuroblasts revealed by single-cell RNA sequencing. Nat. Commun. 13, 1247. 10.1038/s41467-022-28915-3.

