## Supplementary Figures for "The microcephaly protein Abnormal Spindle has an essential role in symmetrically dividing neural precursors to promote brain growth and development"

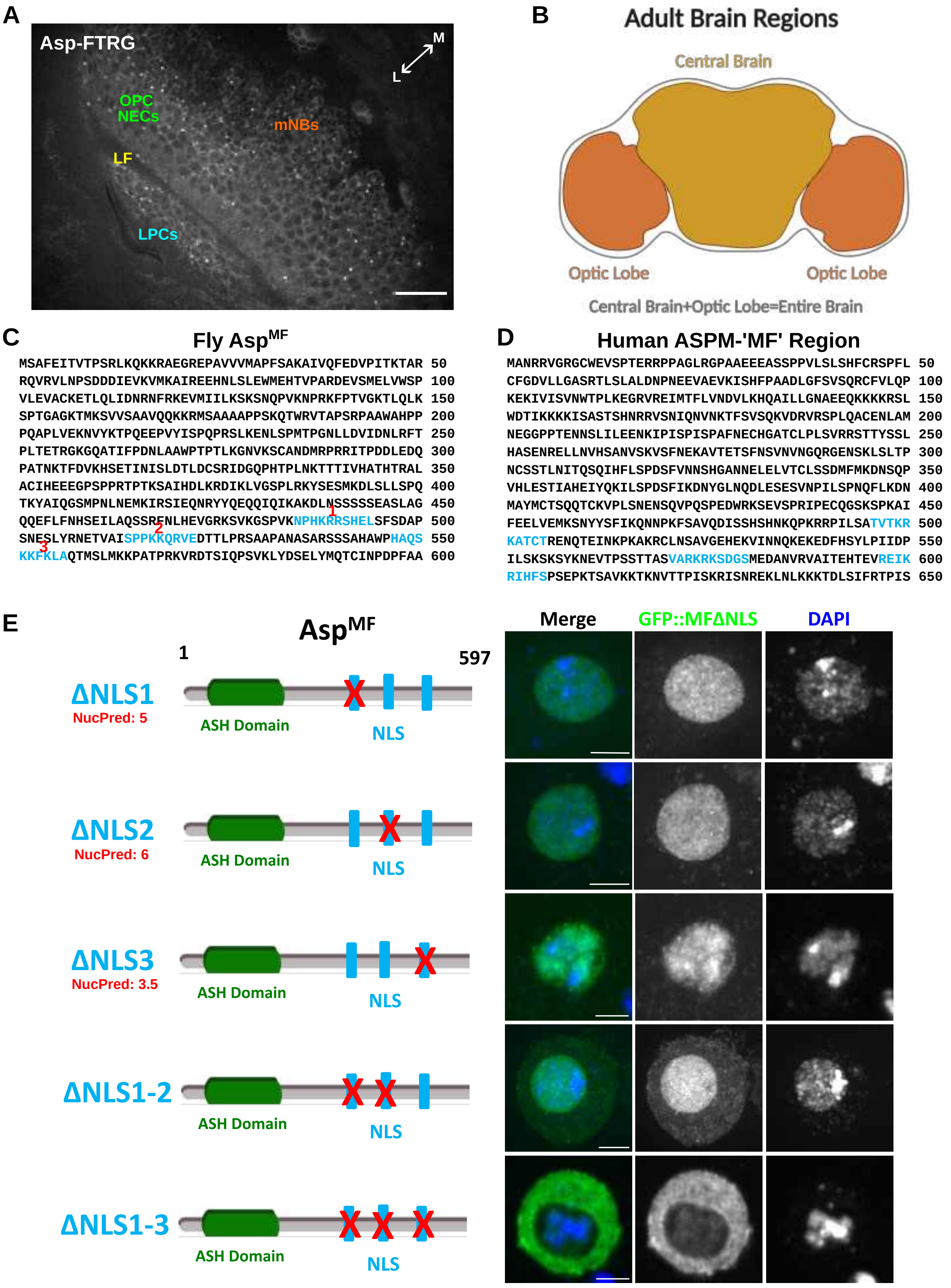


### Supplementary Figure 1 Chakraborty

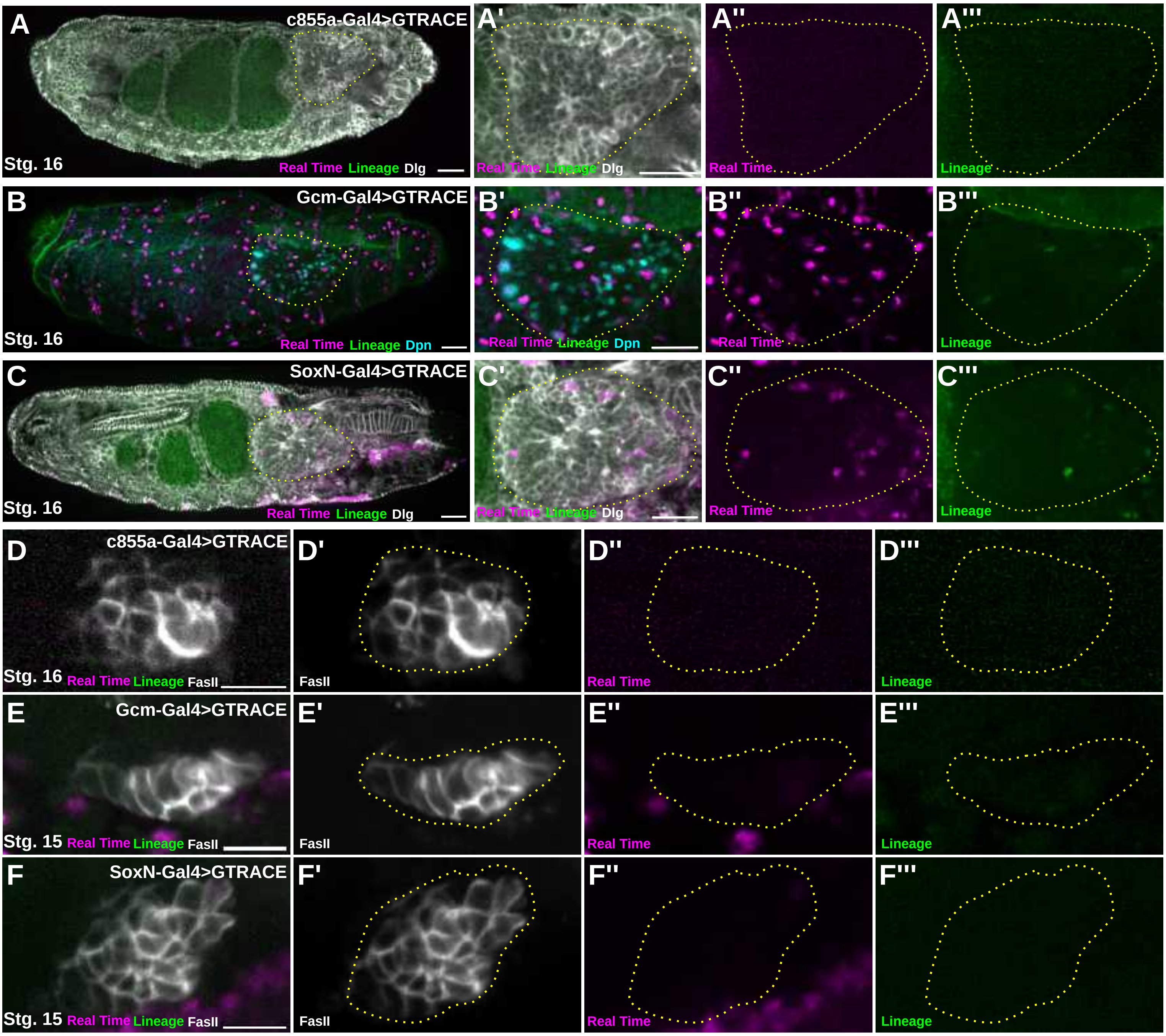


**Supplementary Figure 2 Chakraborty**

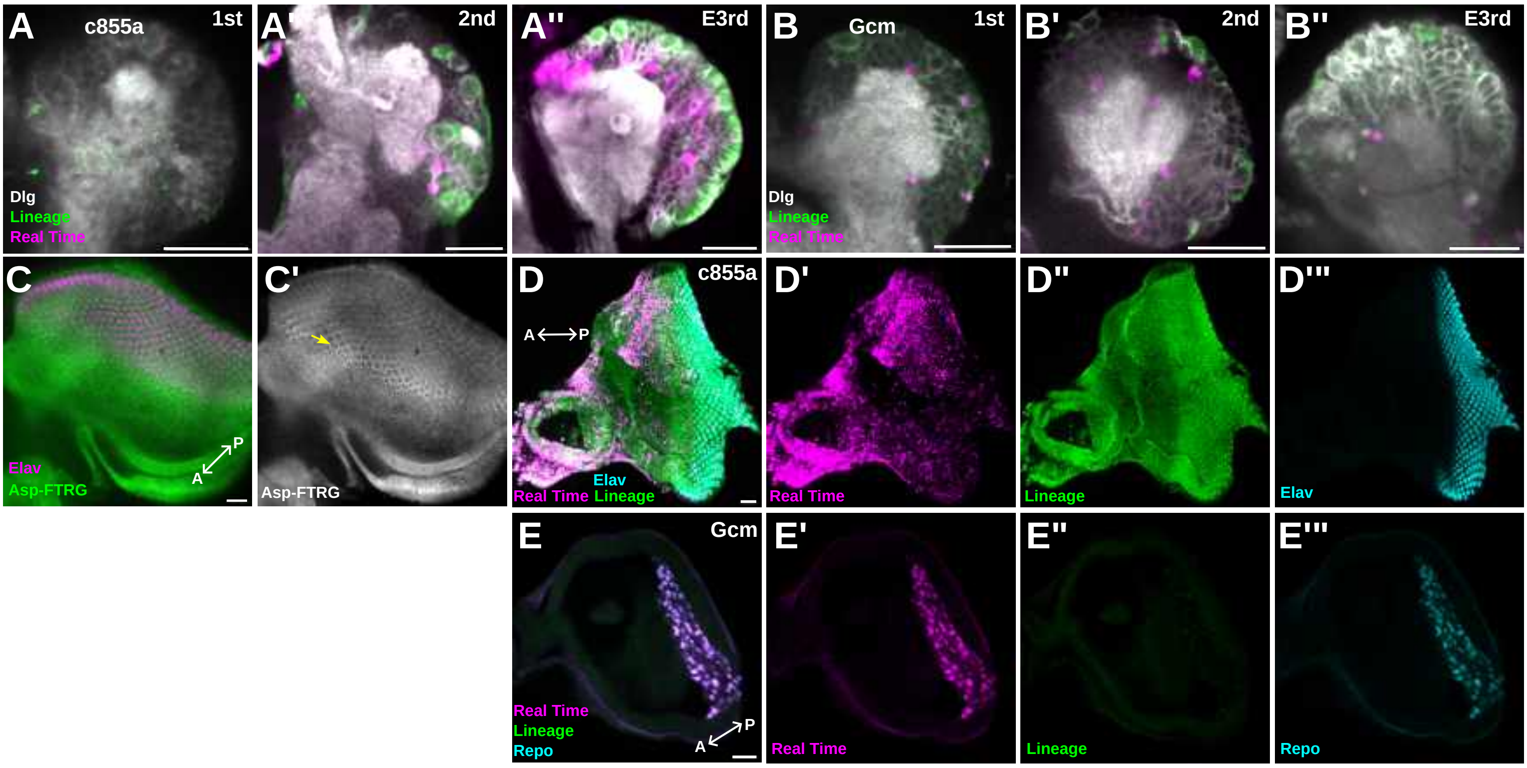


### Supplementary Figure 3 Chakraborty

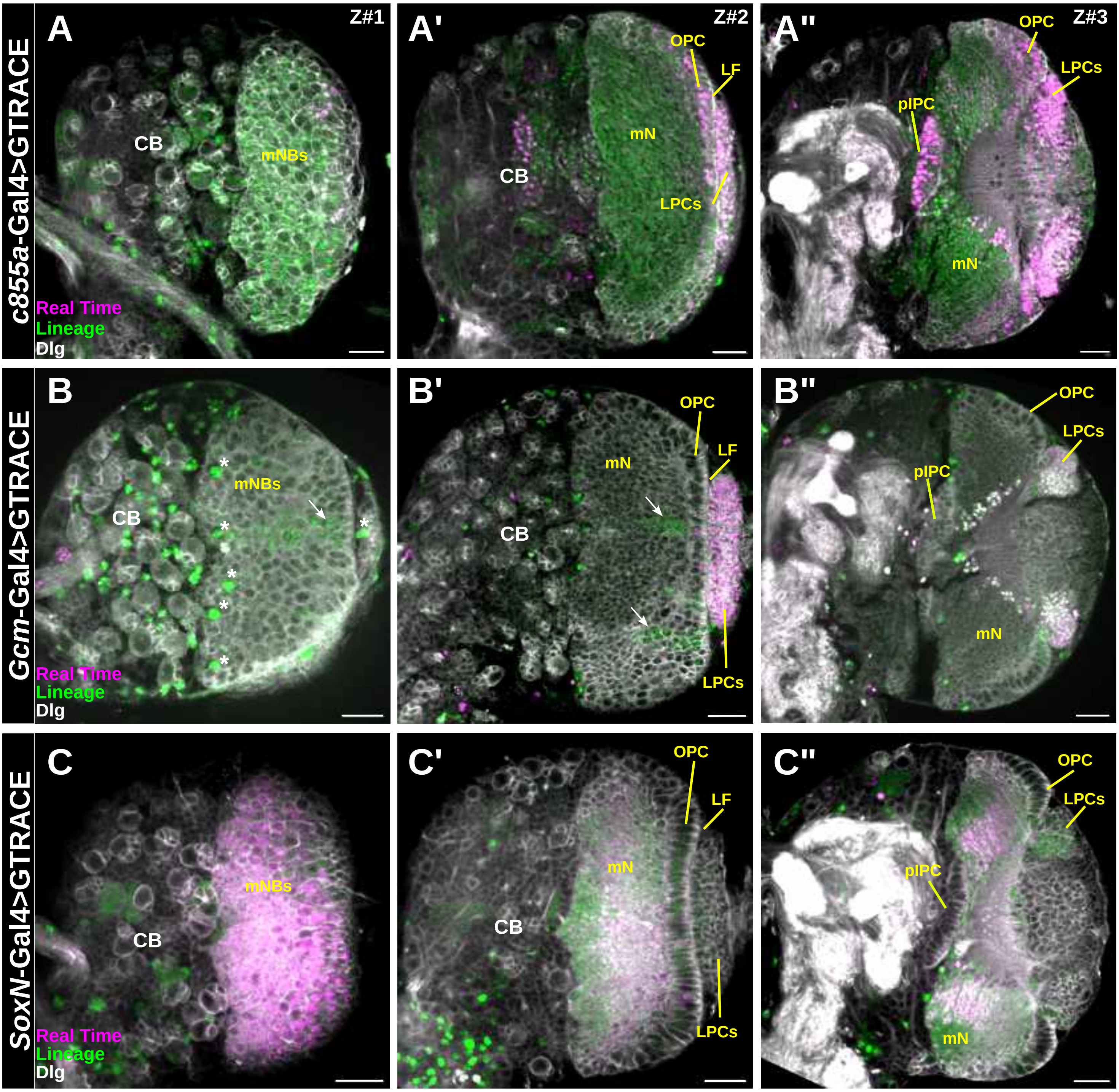


**Supplementary Figure 4 Chakraborty**

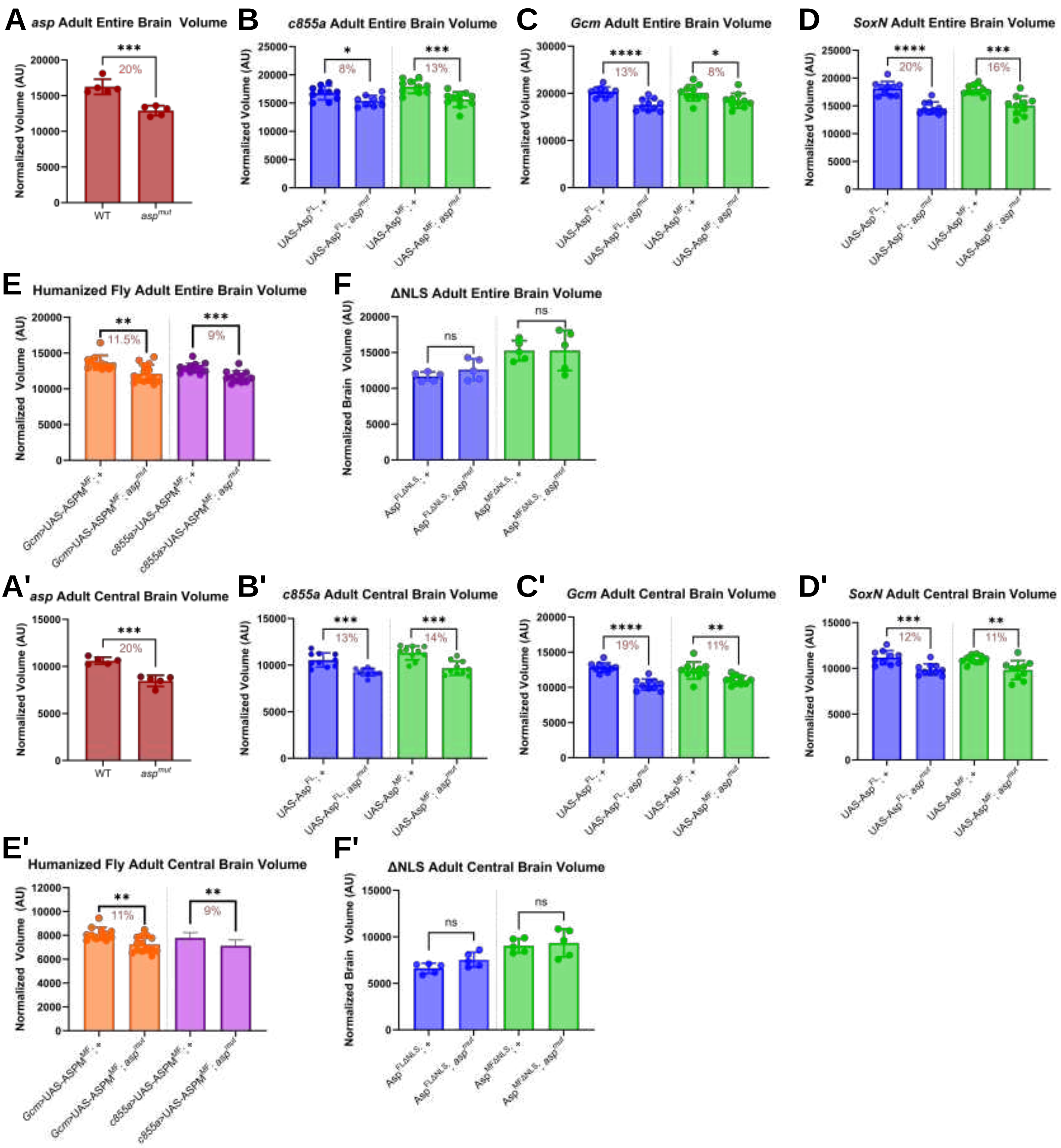


**Supplementary Figure 5 Chakraborty**

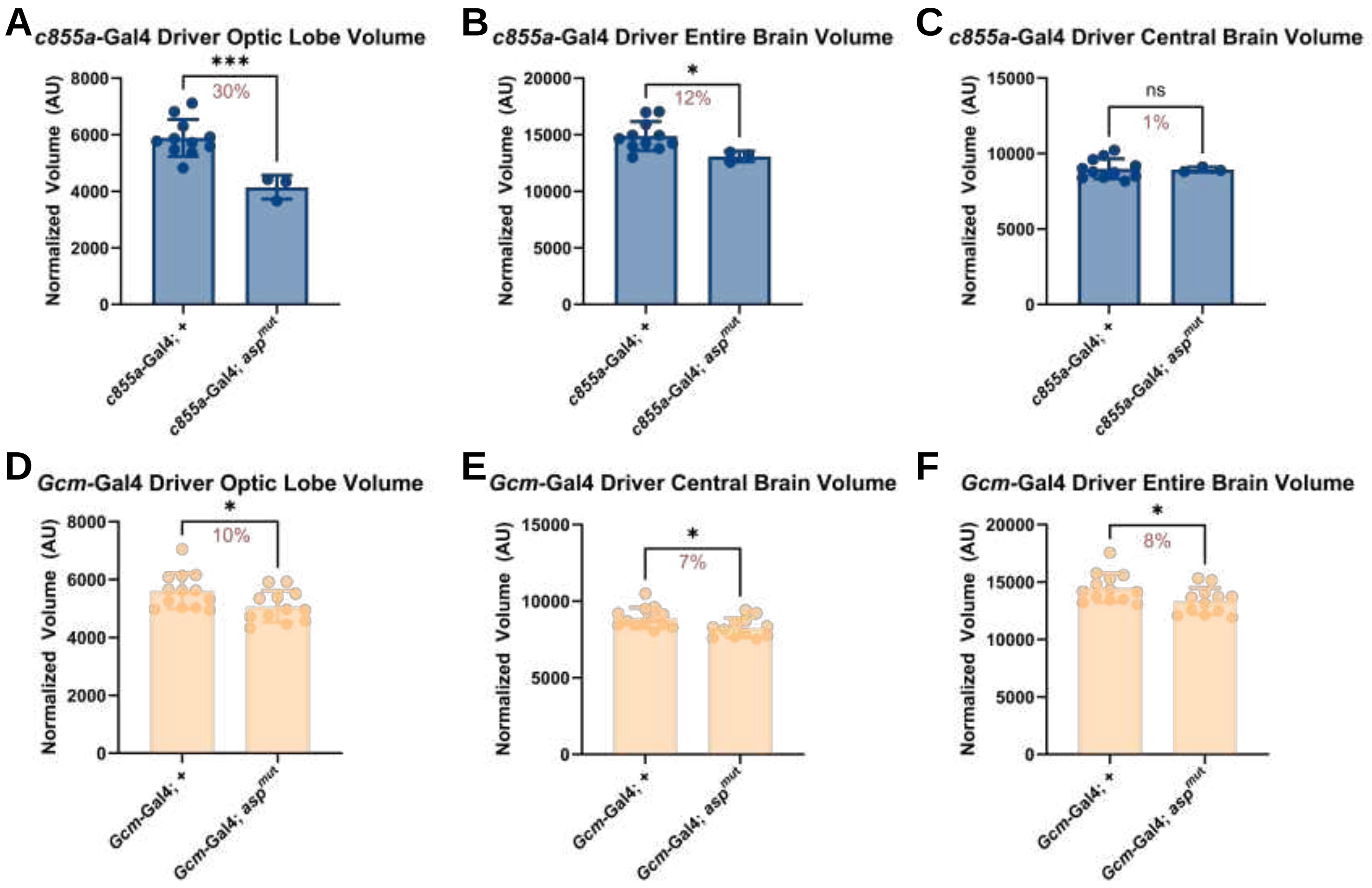


**Supplementary Figure 6 Chakraborty**

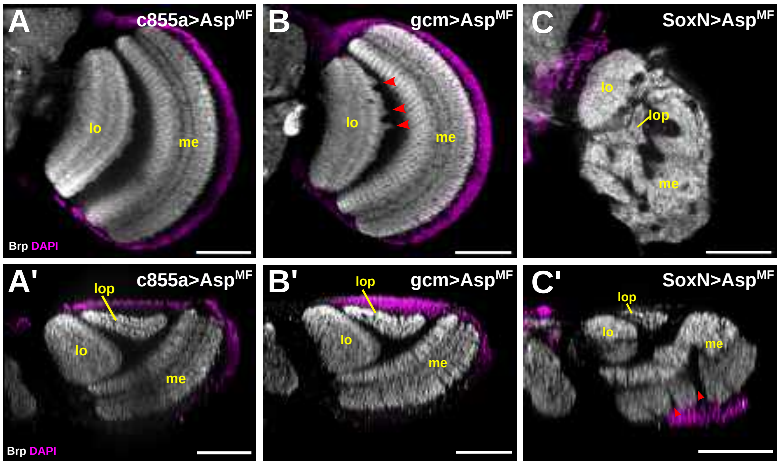


### Supplementary Figure 7 Chakraborty
